# Consistent marine biogeographic boundaries across the tree of life despite centuries of human impacts

**DOI:** 10.1101/2020.06.24.169110

**Authors:** Luke E. Holman, Mark de Bruyn, Simon Creer, Gary Carvalho, Julie Robidart, Marc Rius

## Abstract

Over millennia, ecological and evolutionary mechanisms have shaped macroecological distributions across the tree of life. Research describing patterns of regional and global biogeography has traditionally focussed on the study of conspicuous species. Consequently, there is limited understanding of cross-phyla biogeographic structuring, and an escalating need to understand the macroecology of both microscopic and macroscopic organisms. Here we used environmental DNA (eDNA) metabarcoding to explore the biodiversity of marine metazoans, micro-eukaryotes and prokaryotes along an extensive and highly heterogeneous coastline. Our results showed remarkably consistent biogeographic structure across the kingdoms of life, which were underpinned by environmental and anthropogenic influence. Additionally, metazoan communities displayed biographic patterns that suggest regional biotic homogenisation of conspicuous species. Against the backdrop of global pervasive anthropogenic environmental change, our work highlights the importance of considering multiple domains of life to understand the maintenance and drivers of marine biodiversity across broad taxonomic, ecological and geographical scales.

## Introduction

Researchers have long recognised the importance of grouping global biota into distinct, geographically separated regions. Delineating these biogeographic areas is important to understand the factors shaping the range limits of species^1^, to designate key geographic areas for biodiversity conservation^2^ and in formulating predictive responses to environmental change^3,4^. One of the first efforts to define geographic regions of terrestrial biota were Alfred Russel Wallace’s so-called ‘Zoological Regions’^5^, which included six major regions (hereafter realms) that are still recognised in subsequent classifications today^6^. The drivers responsible for these geographic classifications are frequently environmental conditions or physical barriers. Biogeographic studies have shown that deep divergence in the geographic arrangement of terrestrial biota arose as a result of plate tectonics, while shallow divergence has been most frequently attributed to climatic conditions^7^. In aquatic ecosystems, the relative importance of macroecological drivers is less understood, although both climatic (e.g. temperature)^8^ and tectonic forces^9^ have been identified as key determinants of marine biogeographic patterns. Recent studies have successfully partitioned the oceans into distinct ecoregions (i.e. a geographically defined area, smaller than realm, that contains characteristic assemblages of species)^1,10^, but the description of marine ecoregions has mostly considered conspicuous or well-described species. Similarly, most macroecological marine research has focussed on readily identifiable eukaryotic species, principally metazoans^11^, although demonstrable progress has been made in understanding global patterns of marine microbes^12^. In line with recent studies demonstrating strong cross-phylum interdependence^13^, there is an increasing need to include prokaryotic species in our assessment of biogeographic patterns. The language of macroecology and microbial ecology is similar, both examining the incidence of species across different spatial scales, but these fields have long progressed independently. As a result, relatively few studies have explored biogeographic patterns simultaneously for both microscopic and macroscopic life, with examples of consistent and inconsistent patterns across different taxa^13-15^. Work is thus needed to explore the consistency of biogeographic breaks across different kingdoms of life.

Human driven habitat destruction, pollution and the introduction of non-native species are the main drivers of recent global biodiversity change^3^ and therefore have the potential to alter geographic patterns of biota at multiple spatial scales. Cumulatively, anthropogenic stressors not only threaten vulnerable native species but whole-community structure and function^3,16,17^. The magnitude and direction of human impacts are complex, with evidence for both gains and losses in local species richness across biomes^18-20^. However, a consistent global pattern is emerging, with a recent and rapid increase in species turnover^19^ and an associated increase in community similarity (β diversity) between two or more geographically separated sites^20^. Incidences of increased community similarity are known as biotic homogenisation^21^ and are driven by human activities that promote extinctions and the introductions of non-native species. In light of growing evidence that taxonomic, phylogenetic and functional diversity are strongly correlated^22^, the homogenisation of biological communities has the potential to negatively affect ecosystem function. Furthermore, even uncommon species within an ecological community can contribute significantly to ecosystem function^23^, demonstrating the importance of studying inconspicuous species to preserve ecosystem health. Studies have shown evidence for biotic homogenisation around the globe, with examples from plants^18^, vertebrates^24^ and invertebrates^25^ demonstrating alteration of terrestrial biogeographic patterns. However, many studies are of limited taxonomic scope, focussing on highly conspicuous species for which reliable data can be relatively easily produced^24,25^. Thus, most work overlooks inconspicuous species (e.g. microbes and microscopic eukaryotes), which show vastly different replication, demographic and dispersal patterns compared to metazoans^13^, but are known to be key actors shaping the assembly of ecological communities and ultimately underpin ecosystem functioning^26^. Taken together, a more comprehensive characterisation of ecological communities is clearly needed when testing the role of biotic homogenisation on biogeographic patterns.

The advent of high-throughput sequencing has revolutionised our understanding of microbial life, with studies examining global patterns of prokaryotic life now increasingly common^12^. Moreover, the recent and rapid development of methods to infer the incidence of larger organisms using genetic material isolated from environmental samples (known as environmental DNA or eDNA) has provided an unparalleled ability to identify species across the entire tree of life^14,15,27^. Together, these methods can rapidly generate standardised biodiversity data for entire communities at an unprecedented resolution, thereby minimising regional and taxonomic biases. Such studies provide datasets that can be analysed without taxonomic assignment and DNA samples can be repurposed to test novel hypotheses. A common technique is to amplify DNA barcodes from eDNA and use high-throughput sequencing to produce high-resolution biodiversity data. This method (eDNA metabarcoding) has been shown to reliably detect organisms across many different ecosystems^27^, but has infrequently been applied to understand spatial patterns of biodiversity across different kingdoms of life^12,15^.

A unique geographic setting for testing biogeographic hypotheses is the South African coastline, where two large water masses (the Atlantic and Indian Oceans) meet, and a wide variety of abiotic and biotic conditions are found in a relatively small region. This coastline has three well-defined coastal ecoregions bounded by the cold western boundary Benguela Current and the warm oligotrophic eastern boundary Agulhas Current. These ecoregions have been established on the basis of studies over several decades involving a number of conspicuous metazoan taxa^2,28^. Additionally, there is evidence for human exploitation of marine resources in the region spanning thousands of years^29,30^ and some areas of the coastline have been subject to heavy maritime activity for centuries^31^. Other human activities also prevail such as the establishment of aquaculture facilities or the construction of harbours and breakwaters^29,30^. Thus, the South African coastline is an ideal study system to explore the mechanisms shaping biogeographic boundaries.

Here we used eDNA metabarcoding to compare the biogeography of multiple marine kingdoms along the diverse coastline of South Africa and to test the effects of life-history strategies and anthropogenic and natural pressures on biogeographic patterns. We first investigated the consistency of biogeographic boundaries across metazoans, protists and bacteria, and to what extent these patterns could be explained by environmental conditions. We then evaluated if there was evidence for homogenisation of ecological communities along a coastline that has been affected by centuries of human impacts, and to what extent these patterns persisted across taxonomic groups.

## Methods

### Field sampling

We sampled a range of sites along 2,000 km of coastline (Fig. 1) between October and November of 2017 (see details in Supplementary Table 1), covering the three major marine ecoregions of South Africa. In order to assess the effects of anthropogenic impacts, we compared human altered ‘artificial’ sites (e.g. recreational marinas, harbours) and relatively unaltered rocky shore and natural harbour sites, hereafter ‘natural’ (see Supplementary Table 1). The artificial sites were previously sampled in Rius, et al. ^32^, and six adjacent natural sites were selected. The natural sites were the nearest non-developed sites with matching aspect and exposure (Fig. 1) to each of the artificial sampling sites. Three 400 ml seawater samples were filtered with 0.22 μm polyethersulfone membrane Sterivex filters (Merck Millipore, MA, USA) following the sampling scheme of Holman, et al. ^33^ at each sampling site. Consequently, we sampled a total of 1,200 ml of seawater per site, a volume that has been shown to differentiate fine scale (<1 km^2^) community structure in marine systems^33,34^. Filters were immediately preserved at ambient temperature with the addition of 1.5 ml of Longmire’s Solution for preservation until DNA extraction. Field control filters and equipment cleaning blanks were taken, transported, stored and sequenced as the rest of the field samples.

**Figure 1.**
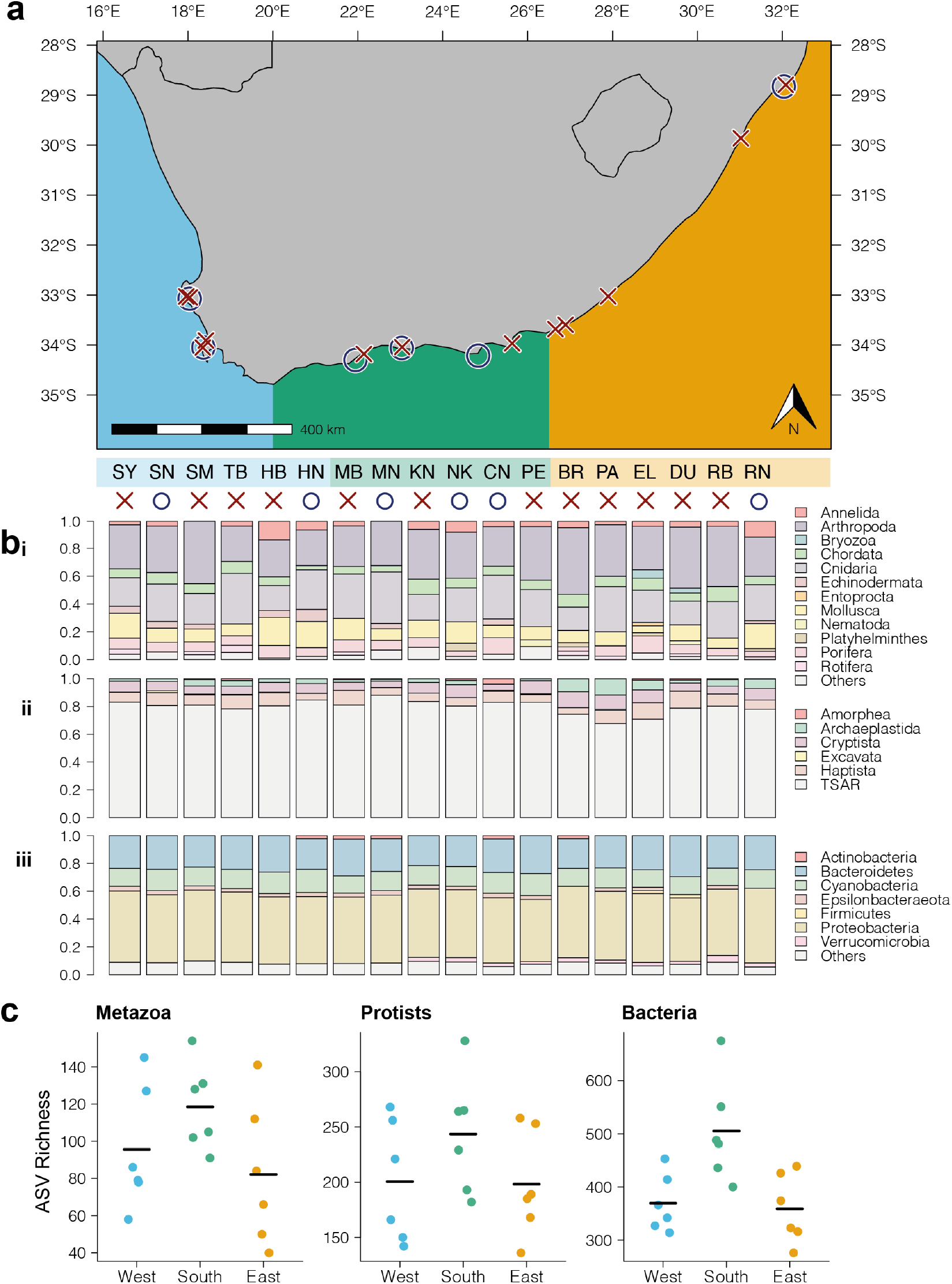
**a** Map of South Africa indicating the sampling sites and the site types (red crosses are artificial sites and blue circles natural sites). Site codes as in Supplementary Table 1; **b** Proportion of ASVs per phyla across each site for **i** metazoans, **ii** protists and **iii** bacteria. Each bar represents a site indicated by the site code as in Supplementary Table 1; **c** Amplicon sequence variant (ASV) richness per site separated by coast and taxonomic group, black line indicates mean ASV richness.

### Environmental DNA extraction

We used a PCR-free laboratory separated from the main molecular biology laboratory facilities. No post-PCR or high concentration DNA samples were permitted in the laboratory. All surfaces and lab equipment were cleaned thoroughly before use with 1.25% sodium hypochlorite solution (3:1 dilution of household bleach). DNA extraction followed the SX^CAPSULE^ method from Spens, et al. ^35^. Briefly, filters were first externally cleaned with sterile water and Longmire’s Solution was removed from the filter outlet using a sterile syringe, 720 μl Buffer ATL (Qiagen, Hilden, Germany) and 80 μl Proteinase K (20mg/ml) was added and filters were incubated overnight at 56°C. The lysate was then removed from the filter inlet and subjected to DNA extraction using the Qiagen DNeasy Blood and Tissue Kit under the manufacturers recommended protocol. DNA was eluted using 200 μl Qiagen Buffer AE and re-eluted once to increase DNA yield. All DNA samples were checked for PCR inhibition using the Primer Design Internal Positive Control qPCR Kit (Primer Design, Southampton, UK) with 10 μl reactions under the manufacturer recommended protocol. Inhibition was detected by an increase of >1.0 Ct in reactions containing eDNA compared to reactions with extraction controls. As inhibition was detected in a minority of samples, all samples were treated using the Zymo OneStep PCR Inhibition Removal Kit (Zymo Research, California, USA) following the manufacturer recommended protocol. Inhibited samples showed no evidence for inhibition post cleaning.

### High throughput eDNA amplicon sequencing

Different sets of primers were used to generate three separate eDNA metabarcoding libraries for all samples. Two gene regions were selected to target broad metazoan/eukaryotic diversity, a section of the V4 region of the nuclear small subunit ribosomal DNA (hereafter 18S)^36^ and a section of the standard DNA barcoding region of cytochrome c oxidase subunit I (hereafter COI)^37^. One gene region was used to target prokaryotes, the V3-V4 hypervariable region of prokaryotic small subunit ribosomal DNA (hereafter 16S)^38^. Illumina unique double-indexed metabarcoding amplicon libraries were constructed with a two-step PCR protocol as detailed in Holman, et al. ^33^. The first PCR setup was performed in a PCR-free laboratory. The three eDNA samples per site were pooled and three independent technical replicates were sequenced per pool. The process per sequenced pool was as follows. The first PCR reaction was conducted in triplicate in a total reaction volume of 20 μl. Each reaction contained 10 μl Amplitaq GOLD 360 2X Mastermix (Applied Biosystems, California, USA), 0.8 μl (5 nmol ml^-1^) of each forward and reverse primers and 2 μl of undiluted environmental DNA template. The reaction conditions for PCR were an initial denaturation step at 95°C for 10 minutes followed by 20 cycles of 95°C for 30 seconds, variable annealing temp (46°C for COI, 50°C for 18S and 55°C for 16S) for 30 seconds, and extension at 72°C for 1 minute. A final extension at 72°C was performed for 10 minutes. The triplicate first PCR replicates were then pooled and cleaned using AMPure XP beads (Beckman Coulter, California, USA) at 0.8 beads:sample volume ratio following manufacturer’s instructions. The second PCR reaction was conducted in a total volume of 20 μl containing 10 μl Amplitaq GOLD 360 2X Mastermix, 0.5 μl (10 nmol ml^-1^) of both forward and reverse primers and 5 μl of undiluted cleaned PCR product from the first reaction. PCR conditions were an initial denaturation step at 95°C for 10 minutes followed by 15 cycles of 95°C for 30 seconds, annealing at 55°C for 30 seconds, and extension at 72°C for 1 minute. A final extension at 72°C was performed for 10 minutes. PCR 2 products were cleaned using AMPure XP beads as above. Negative control samples for the filters, extraction kit, PCR1 and 2 were included in library building and sequenced alongside experimental samples. Products were quantified following the manufacturer’s instructions using the NEBNext Library Quant qPCR kit (New England Biolabs, Massachusetts, USA) and then normalised and pooled at an equimolar concentration for each marker. Each gene region was sequenced independently using a V3 paired-end 300bp reagent kit on the Illumina MiSeq Instrument with 5% PhiX genomic library added to increase sequence diversity.

### Bioinformatics

Raw sequences were de-multiplexed using the GenerateFastQ (v2.0.0.9) module on the MiSeq control software (v3.0.0.105). Cutadapt (v2.3)^39^ was used to filter sequences to include only those that contained both the forward and reverse primer sequence across both read pairs for each gene fragment, remaining sequences then had the primer region removed for each gene fragment using the default settings. Sequences were denoised using the DADA2 pipeline (v1.12)^40^ in R (v3.6.1)^41^ with the default parameters unless noted as follows. Sequences were filtered to retain only pairs of reads with an expected error of 1 or fewer per read. Read trimming was performed after manual examination of the read quality profile, the forward reads were trimmed to 250bp (COI), 240bp (18S) and 240 bp (16S) and the reverse reads were trimmed to 230bp (COI), 220bp (18S) and 220 bp (16S). As each marker was sequenced separately, the differences in read trimming length reflect typical variation in sequencing runs rather than any biological difference. The error rates per run were estimated and used to perform the denoising using the DADA2 algorithm. The denoised sequence pairs were then merged and resulting sequences were truncated if they were outside of the expected gene fragment range (303-323bp for COI, 400-450bp for 18S and 390-450bp for 16S). Chimeras were identified and removed before assembling a sample by ASV (amplicon sequence variant) table for analysis. The denoised ASVs were then curated using the default settings of the LULU algorithm^42^ which merges sequences based on sequence similarity and co-occurrence. Assigning taxonomy to a set of unknown sequences is a difficult task, particularly considering many marine species lack DNA barcodes, are undescribed, or have erroneous barcodes in online public databases. We therefore focused our analysis at a higher taxonomic level than species, assigning taxonomy to sequences from the COI and 18S data as follows. The RDP classifier (v2.13)^43^ was used to assign taxonomy for COI using a previously published COI database^44^ (v4.0) and a modified version of the SILVA database^45^ (v3.2 from https://github.com/terrimporter/18SClassifier). As species level assignments have been shown to be accurate for COI data^33^ an unconstrained (no limits on sequence similarity or match length) BLAST search (v2.6.0+) was performed for each sequence against the entire National Centre for Biotechnology Information nt database (downloaded on 16^th^ May 2019), 200 hits per sequence were retained (*-num_alignments*). These sequences were then parsed using an R script to exclude hits below 65% coverage, remaining assignments with percent identity above 97% for COI were used to collapse reads for ASVs assigned to the same species. Recent analyses have suggested that only exact (100% identity) matching of sequences to reference data is appropriate for species assignment for the prokaryotic 16S region^46^. The 16S sequences were matched to the SILVA database (release 132)^45^ using the default settings of the *assignTaxonomy* function from the DADA2 package to assign taxonomy at genus level or above. The incidence of NUMTs (nuclear mitochondrial DNA) and chimeras in the final ASV list was evaluated following Supplementary Information 1.

The following quality control filters were applied to the ASV by sample table produced by DADA2. First, the minimum number of reads per observation was set at three. Any ASVs not represented in at least one other sample were discarded. ASVs were then filtered to retain only those found in all three technical replicates. For any ASV found in the negative control samples, the largest value among the control samples was used as the zero value for all other samples (i.e. any smaller values found in non-control samples were set to zero). The COI and 18S datasets were then subset by the RDP classifier taxonomic assignments to produce datasets for the protists and metazoans as follows. Phylum level assignments above a threshold of 30, a value well above that shown to accurately assign phylum level taxonomy ^44^, were parsed to include phyla that contained only metazoan or protist members for each group respectively, other assignments or unknown assignments were discarded. This resulted in a protist and metazoan dataset for each marker, which were subsequently used for separate analyses using these groupings. The 16S data was parsed to include only bacterial ASVs. Within each taxonomic dataset samples were then rarefied to the smallest number of reads (see Supplementary Table 2). Technical replicates were then collapsed to produce a dataset containing the mean value of rarefied reads per ASV. Finally, ASVs assigned using BLAST (with no cases of multiple matches of equal quality) to the same species in the COI dataset were combined by summing reads per site. The taxonomic assignment method used for the 16S data assigns to genus, and species level assignments are not possible using the selected 18S region, so no ASVs from these datasets were collapsed. In order to explore broad scale patterns of taxonomic diversity, the number of ASVs per phyla and number of rarefied reads per phyla were collapsed to produce per site assessments of taxonomic composition. As phylum level phylogeny is not resolved for all protist species, the protist dataset was grouped by supergroup designations according to Burki, et al. ^47^. For plots, phyla represented by less than 2% of ASV counts were concatenated in an ‘other’ category.

### Abiotic, human impact and geographic data

*In situ* temperature data reflects a snapshot of the total conditions experienced across the lifetime of the species that make up marine communities. Therefore, abiotic variables for the sites covering an ecologically relevant timescale were sourced as follows. High resolution (1km^2^) remote sensing average daily sea surface temperature (SST) data derived from multiple satellite missions, combined with *in situ* data^48^ was parsed in R to find the nearest datapoint to each site. For each point, a mean from two years of data from November 2017 was calculated. Interpolated average (2005-2017) sea surface salinity (SSS) data (0.25° grid resolution) generated using gliders, oceanographic casts etc. from the 2018 World Ocean Atlas^49^ was parsed to include only surface data for the sites. Monthly global ocean colour data (4km^2^) derived from multiple satellite missions^50^ was parsed to calculate an average value for chlorophyll *a* density per site across two years from November 2017. Finally, a previously described^51^ 1km^2^ global resolution cumulative index for anthropogenic impact on marine ecosystems, comprising fishing pressure, climate change, shipping and land-based pollution, was parsed to produce a value for each site cumulatively across the entire period for which data were available (2003-2013). These global datasets have excellent temporal resolution, but are only appropriate for testing large-scale patterns as they have limited ability to discriminate highly localised observations.

### Ecological statistics

Differences in the mean number of ASVs per coastline were assessed using an analysis of variance (ANOVA) after testing for normally distributed residuals using a Shapiro-Wilk test and equal variance between coasts using a Bartlett test. A Tukey’s Honest Significant difference test was used to evaluate significant ANOVA results. Differences in community similarity were assessed using a Permutational Multivariate Analysis of Variance (PERMANOVA)^52^ implemented in R with the function *adonis* from the package *vegan* (v2.5-6)^53^ to assess differences in multivariate centroids and dispersion between coastlines. The PERMANOVA was conducted on a matrix of Jaccard dissimilarities as this ecological index has been shown to be appropriate for biogeographical studies^54^. Significant pairwise differences were assessed using the R function *adonis*.*pair* from the *EcolUtils* package (v0.1)^55^. To analyse if groups of samples have a difference in intra-group community variation, also known as heterogeneity of multivariate dispersion, the PERMDISP2 procedure^56^ was used, implemented in the R function *betadisper* from the *vegan* package. The pairwise group differences in heterogeneity of multivariate dispersion in the case of a global significant result from *betadisper* were analysed using a Tukey’s Honest Significant difference test. Non-metric multidimensional scaling ordinations (nMDS) were calculated using Jaccard dissimilarities and the R function *metaMDS* from the *vegan* package.

The influence of the abiotic and human impact data on the observed patterns of beta diversity were evaluated as follows. It has previously been common to use a partial Mantel test to evaluate the effect of a distance matrix (frequently environmental variables) on a second distance matrix (species composition) while ‘cancelling out’ the effect of a third matrix (geographic distance). However, this approach has been shown to be sensitive to spatial autocorrelation common in ecological datasets^57^. A recently developed method^57^, which corrects spurious inflations of the parameter estimate for Mantel tests, was implemented in R. Across each taxonomic group Mantel tests were conducted comparing Jaccard dissimilarity against Euclidean distance for each environmental variable. For each test Moran spectral randomisation was performed including the geographic distance data with 10,000 permutations to assess statistical significance using the *msr* function in the *adespatial* package (v0.3-8).

Explanatory variables which had some correlation with the community dissimilarity after adjusting for geographic distance were then evaluated as follows. First, a distance-based redundancy analysis (dbRDA)^58^, regressing site Jaccard dissimilarities against all remaining variables, was performed using the R function *dbrda* from the *vegan* package. The significance of terms was assessed with 10,000 permutations. The dbRDA ordination allows us to examine linear changes in the beta diversity in response to a number of predictor variables in tandem, and also to explore their relative impact. The function *varpart* from the *vegan* package was then used to partition the variance in the community dissimilarity by the environmental variables. We then used a generalised additive model to visualise the variation of each significant variable across the nMDS space via a restricted maximum likelihood 2D smoother, implemented in the function *ordisurf* in the R package *vegan*.

Distance-decay relationships were explored by first measuring compositional similarity (1-Jaccard index) for each pair of sites, and then calculating distances between pairs of sites by drawing a continuous transect 1km offshore parallel to the high-water mark using Google Earth Pro (v 7.3.2.5776), taking the distance along the transect to measure distance between sites. We then used these data in least-square regression models using the R function *lm* with an interaction function between distance and site type (artificial or natural) terms against compositional similarity as a response term. The compositional similarity values were log10 transformed to linearise the response, untransformed values of zero (no overlap of species) were omitted to avoid infinite response variable values.

## Results

### Sequencing

A total of 66.25 million sequences were produced across the three sequencing runs. The number of unfiltered raw reads per experimental sample ranged from 61,958 to 859,580, with an average per sample across all three markers of 347,536 ± 109,665 (*s*.*d*.) (see Supplementary Table 2 for further details). Negative control samples exhibited very low levels of cross-contamination (Supplementary Information 2).

### Taxonomic assignments & alpha diversity

Analyses for each taxonomic group are presented for the marker which had the largest number of observations (COI for metazoans, 18S for protists), analyses for the remaining subsets are presented in Supplementary Information and were consistent with the main results. After taxonomic assignment to phyla 1,054, 1,433 and 2,826 ASVs were retained for the metazoans, protist and bacteria datasets respectively. Across all taxonomically grouped datasets the majority of detected ASVs came from a small number (<5) of phyla or supergroups (Figure 1b). This pattern was consistent across sites and no major changes in the identity of ASVs at phyla or supergroup level across the study region were detected (Figure 1b).

Across all markers the greatest mean ASV richness was found along the southern coast (Fig. 1). However, a one-way ANOVA showed a significant difference (F2,15=7.18, p=0.007) between coastlines only in the bacterial dataset with no difference found in the metazoan (F2,15=1.941, p=0.178) or protist data (F2,15=1.416, p=0.273). A post-hoc Tukey test of the bacterial data (Supplementary Table 3) showed that the east and west coasts had significantly fewer ASVs compared to the south coast, but that they were not significantly different to one another in overall ASV richness.

### Beta diversity

Across all three taxonomic groups, non-metric multidimensional ordinations showed clustering of sites consistent with ecoregions previously described in conspicuous metazoan species (Fig. 2). Furthermore, PERMANOVA models showed a significant (p<0.001) effect of coastline in all cases (see Supplementary Table 4 for model output), with pairwise significant differences (p<0.01) between all pairs of coastlines in all taxa (Supplementary Table 5). There was evidence (ANOVA on *betadispr* F2,15=4.09, p=0.038) for heterogeneity of multivariate dispersion in the bacterial dataset (Supplementary Table 6). A Tukey test revealed a significant (p=0.031) pairwise difference between the east and west coast only, in line with the observations of Fig. 2., indicating that for bacteria, sites on the west coast were more variable in community composition than the more homogenous communities found at sites on the east coast. When the datasets were divided by phylum, they demonstrated consistent evidence for the same significant difference between ecoregions across all tested phylum (p<0.05 in all cases, see Supplementary Information 3 for full model output and visualisation). A power analysis (Supplementary Information 4) indicated that the number of ASVs allocated to each phylum was sufficient to detect a significant difference given the study design.

**Figure 2.**
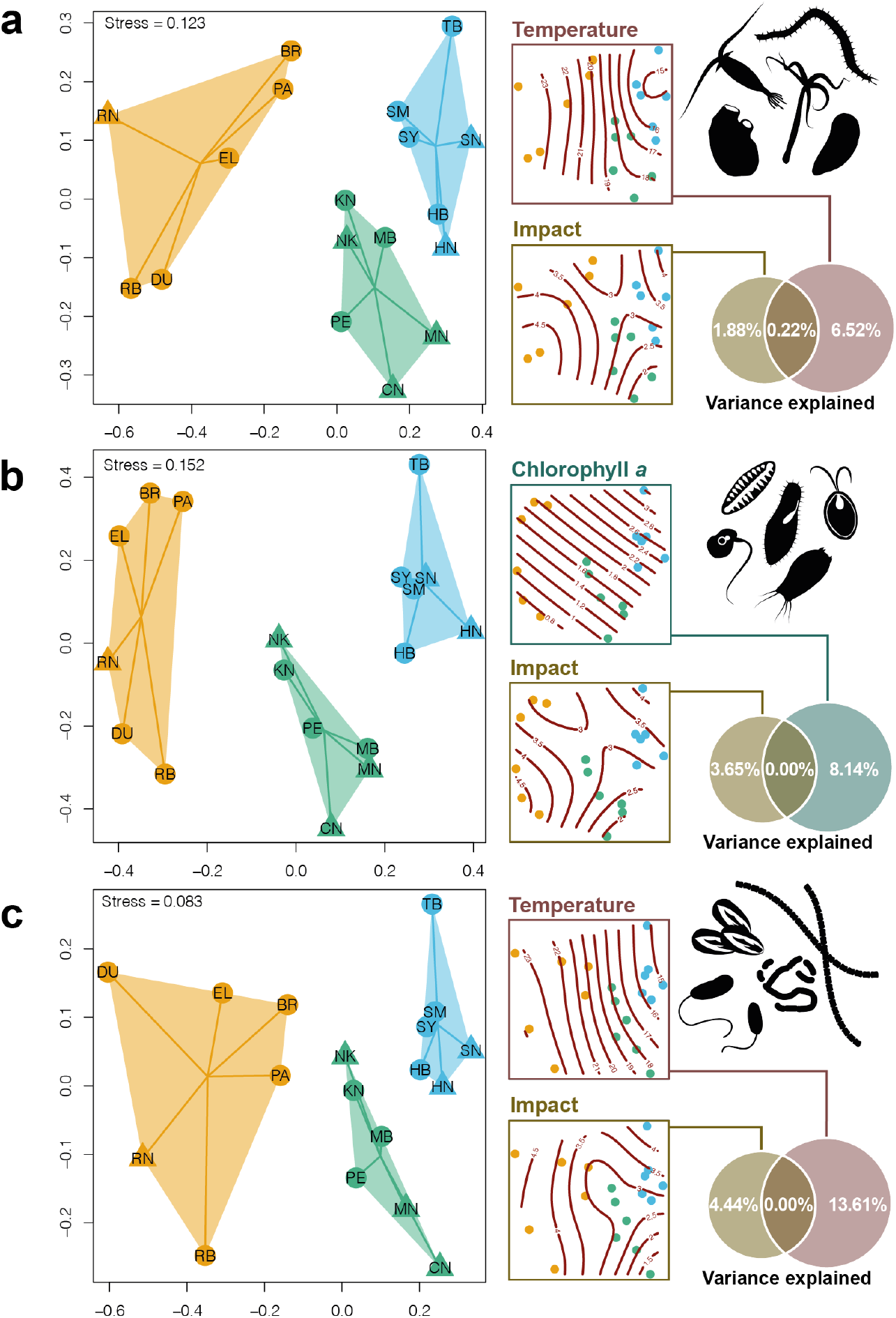
Observed patterns of β-diversity from environmental DNA metabarcoding of: **a** metazoans, **b** protists and **c** bacteria; based on Jaccard dissimilarities between amplicon sequence variants along the coast of South Africa. The first column of plots shows non-metric multidimensional scaling (nMDS) ordinations. Coloured hulls show the spread of the data and lines indicate the spread around the centroid grouped by coast with the east, south and west coasts denoted by orange, green and blue respectively. Site abbreviations correspond with Figure 1, natural sites are denoted with triangles and artificial sites with filled circles. The second column of plots shows the same nMDS ordinations as the first column including the output of a generalised additive model with a 2D smoothed function for each of the significant environmental / impact variables overlaid; temperature – mean sea surface temperature (°C); Chlorophyll *a* – chlorophyll *a* concentration (mg m^-3^); impact – human marine impact score (unitless measurement, see details in text) against the two nMDS axes. The Venn diagrams indicate the percentage total of the variance in the community dissimilarity explained by each significant variable, derived using variance partitioning of a distance-based redundancy analysis.

The corrected Mantel tests indicated that in the metazoan and bacterial datasets, SST and the human impact index were significantly correlated with the observed ASV dissimilarities after geographic distance between sites was accounted for (SST p<0.05 in all cases; human impact p<0.01 in all cases; full model outputs shown in Supplementary Table 7). In the protist dataset chlorophyll *a* concentration and the impact index remained significant (p>0.05 in both cases). In contrast, SSS showed no correlation (p>0.05 in all cases) with observed ASV dissimilarities in any marker. These results indicate that both geographic and environmental distance have some effect on the observed community structure and confirmed that the appropriate variables were retained for analysis in each dataset. In all cases, partial Mantel tests gave similar R statistics and p values (Supplementary Table 7).

The dbRDA showed a significant effect of both environmental variables (p<0.001 for SST and chlorophyll *a* in all cases) and human impact (p<0.05 in all cases) on the site similarity in metazoans, protists and bacteria, full model outputs are presented in Supplementary Table 8. Variance partitioning of the dbRDA models showed that human impact had a relatively smaller contribution to the observed dissimilarities compared to the chlorophyll *a* concentration or sea surface temperature as shown in Figure 2. Across taxonomic groups there was negligible overlap in the variance explained by impact and chlorophyll or temperature. Generalised additive models with a 2D smoothed function showed significant terms (p>0.001) across all models (individual full model outputs shown in Supplementary Table 9), indicating that each variable separately explains variation in the eDNA data between communities. SST and chlorophyll *a* concentration showed surfaces across nMDS plots for all markers (Fig. 2) that were simple, with gradients that were consistent with ecoregions. In contrast, human impact scores showed more complex surfaces with multiple peaks across the ecoregions.

### Distance-decay

Distance-decay slopes for all observations showed an exponential decrease in compositional similarity as the distance between sites increased (Fig. 3a). Regression models of log10 transformed compositional similarity indicated that this slope was statistically significant in all cases (P<0.001 for all taxonomic groups, full model output in Supplementary Information 5). In the metazoan dataset the model showed a significant difference in the slope between artificial and natural sites (F3,76=47.73, p < 0.001). No statistically significant difference was found between site types in the protist or bacteria data, and the same pattern was observed within taxonomic groups for the metazoan and protistan subsets across both the COI and 18S data (Supplementary Information 5).

**Figure 3.**
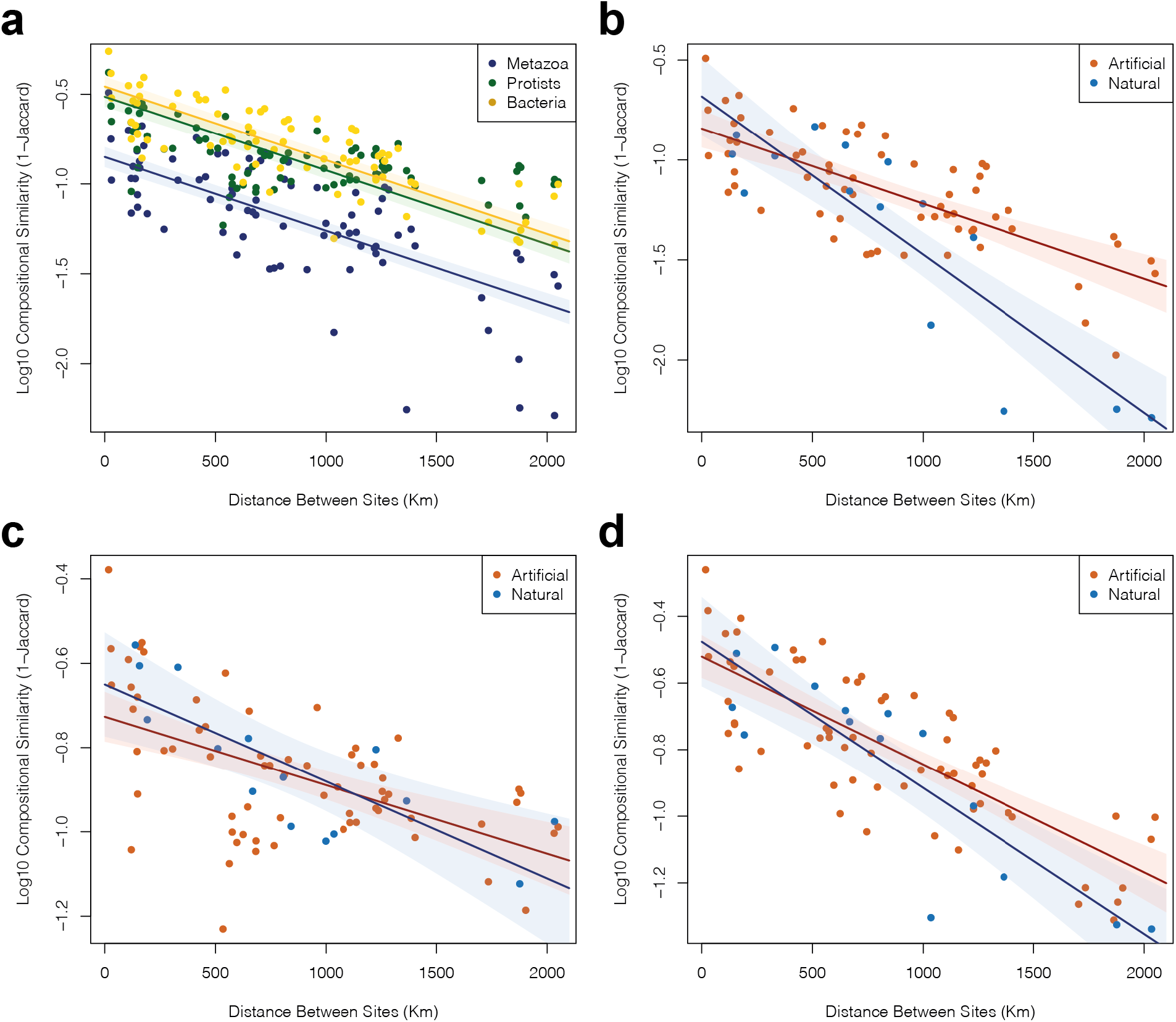
Plots showing distance between sites and community similarity measured using environmental DNA metabarcoding across South Africa. Logarithmically (base 10) transformed compositional similarity against distance is shown in **a**, which includes all datasets. Comparisons between artificial (coloured in red) and natural (coloured blue) sites are shown for **b** metazoans, **c** COI protists and **d** bacteria. 95% confidence intervals from the regression models are shown as light shaded areas around each regression slope.

## Discussion

Here, we showed that metazoans, protists and bacteria can have similar biogeographic patterns along an extensive and heterogenous coastline. We found that these remarkably consistent patterns could be partially explained by measured environmental conditions (chlorophyll *a* and temperature), and to a lesser extent, cumulative human impact. Additionally, we found evidence for anthropogenically driven homogenisation of communities exclusively in metazoans. Collectively, we provide evidence of congruent biogeographic boundaries across vastly different forms of life, and demonstrate that underlying processes, such as anthropogenic alterations, affect biogeographic patterns differentially across taxa.

Prokaryotes and eukaryotes diverged billions of years ago and have since evolved to inhabit a vast range of ecological niches. Previous studies have shown both similar^59,60^ and dissimilar^61^ patterns of β diversity between macro- and microscopic species across environmental and geographic gradients. Recent work has explored biogeographic regionalisation in marine plankton across kingdoms^15^, showing that smaller planktonic organisms such as bacteria may have greater biogeographic structuring compared to larger metazoans or protists^15^. Together this evidence suggests that different ecological processes drive a number of taxon-specific responses to produce patterns that are not universal across systems at different spatial scales^13^. Here we observed similar biogeographic patterns across kingdoms (Fig. 2), providing clear evidence of cross-phyla biogeographical congruence.

Our analyses suggest that environmental variables such as temperature or chlorophyll *a* concentration influenced the structure of marine communities across the study region (Fig. 2). Global studies of biogeographic patterns have shown a central role of temperature in the structuring of both microbial^12^ and larger planktonic life^15,62^ across the ocean. There is growing evidence that the range boundaries of marine organisms closely track their thermal limits^4^. Therefore, a general expectation was that species would remain within their thermal niche resulting in temperature-structured communities as observed here. In contrast to temperature, salinity had a minor role in structuring the studied communities (Fig. 2), an observation previously reported for the global ocean^12^, with exceptions found in microbial^63^ and meiofaunal^64^ life in regions with unusually strong salinity gradients (e.g. Baltic Sea). The SSS range across our study system was very narrow (35.04 - 35.38ppt) and so the negligible observed effect was expected. In the case of protists, biogeographic patterns showed a stronger association with primary productivity (measured here as chlorophyll *a* concentration). Previous research has shown little or no role of productivity in driving coastal and oceanic scale biodiversity patterns^15,62^. However, these studies explored the global role of various environmental variables; the significance of more localised oceanographic systems such as upwelling (as along the western coast of southern Africa) might not be as apparent in global analyses.

Anthropogenic activities are known to alter both the physico-chemical properties of the marine environment and the trophic and ecological properties of ecosystems^65^. In our study system, human impact provided some explanatory power to understand the observed community structure, but to a much lesser extent compared to environmental variables (Fig. 2). The human impact index used here^51^ covered a large number of different types of impact (e.g. pollution, shipping intensity) but even this granular approach adequately explained a small proportion of the total variation in ASVs observed among sampling sites. Previous work on marine metazoans has shown a strong effect of proximate urbanisation^34^ and the ecological drivers produced through anthropogenic activities are well documented^65^. Interestingly, the pervasive and conspicuous urbanisation of the marine environment in the study area showed a much weaker effect on biogeographic patterns than other explanatory variables (Fig. 2). Anthropogenic pressures have become a major ecological driver only relatively recently in evolutionary time, with the most dramatic changes in biodiversity occurring within the 21^st^ century^3^. It is clear that human activities are altering evolutionary trajectories^65^, either through extinction, range expansions or contractions. However, our data suggests that centuries of human impacts in our study system have not yet demonstrably altered the main observed biogeographic boundaries across taxa.

Previous work on biotic homogenisation has shown a dramatic effect on whole communities at both regional^18,66^ and global scales^24,25^. Here, we found evidence for biotic homogenisation along the South African coastline only in metazoan species, with a difference in the slope of a distance-decay relationship between artificial and natural sites. This pattern was consistent for metazoans across the gene regions considered (Supplementary Information 5), with more homogenous community composition between artificial sites. Pervasive vessel activity in the region^31^, along with evidence that artificial environments are hotspots for metazoan invasions^33^, suggest that introduced metazoans are contributing to homogenisation of coastal communities. Further work should incorporate time series data to explore biotic homogenisation, given the significant but minor role of human impact in structuring ecological communities across the region.

Both environmental parameters and species interactions have a clear-cut effect on marine community structure across kingdoms of life^67^, but the comparative role of deterministic (environmental filtering, niche processes, etc.) and stochastic processes (ecological drift, random extinction, etc.) in explaining the observed patterns remains uncertain. The classical deterministic theory (Baas Becking hypothesis^68^, often summarised as ‘everything is everywhere’) of microbial biogeography postulates that due to vast population sizes and dispersal, microbes are found in all environments and the variation in abiotic conditions selects for those that make up the vast majority of species in each region. This theory ignores neutral processes which have been shown to have a critical role in structuring microbial biogeography across biomes^69^. In line with previous efforts studying deterministic and stochastic processes across taxonomic kingdoms^70,71^, we found that the majority of the observed variation could not be fully explained for both prokaryotic and eukaryotic species (Fig. 2). Indeed, recent biogeographic research in the oceans has provided both theoretical^72^ and empirical^15^ evidence of strong biogeographic patterns driven by both stochastic and deterministic forces, but much of the observed variation between communities remains unexplained. Understanding the comparative roles of different community structuring processes requires a more comprehensive examination of observed variance between communities, the interactions between taxa, and the broader role of the environmental conditions where they live.

Several recent innovations will provide valuable data to help uncover the unexplained variation in community structure. For example, the extraction and analysis of sedimentary ancient DNA allows the reconstruction of high-resolution biodiversity change over time^73^. By rewinding the ecological clock, the approach provides evidence to evaluate the role of deterministic processes in reconstructions of community composition through time relative to documented changes in environmental proxies. In conjunction, the analysis of networks from molecular data, in conjunction with life history parameters, could facilitate the hypothesis testing of species interactions that may further elucidate the role of ecological interactions (e.g. Djurhuus, et al. ^14^) in structuring biogeographic patterns. Finally, very high-resolution multi-spectral remote sensing data (e.g. WorldView-3, <100m) will provide unparalleled insights into the role of environmental forces on the distribution of ecological communities^74^.

## Acknowledgments

LH acknowledges the assistance of Molly Czachur and Thomas Grevesse during field surveys, Sophie von der Heyden for lab consumables and assistance with fieldwork logistics, and both Jamie Hudson and Ivan Haigh for assistance with remote sensing data. LH acknowledges Shirley Parker-Nance and the Elwandle Node of the South African Environmental Observation Network for assistance and in-country logistics. All marina owners and operators are acknowledged for field site access. We acknowledge the Environmental Sequencing Facility from the National Oceanography Centre, Southampton for access and sequencing assistance. We thank the IRIDIS High Performance Computing Facility, and associated support services at the University of Southampton. LH was supported by the Natural Environmental Research Council (grant number NE/L002531/1). The UK Research and Innovation Newton Fund (grant: ES/N013913/1) supported LH’s research stay in South Africa.

## Data Availability

Raw Illumina sequencing data are available from the European Nucleotide Archive under study accession number PRJEB38452, sample specific accessions are provided in Supplementary Table 10. Associated metadata, R scripts and intermediate files are available at the following DOI:10.5281/zenodo.4384644.

## Code Availability

All code used in the current study can be found at the following DOI:10.5281/zenodo.4384644.

