## Supplementary Information for "Consistent marine biogeographic boundaries across the tree of life despite centuries of human impacts"

### Supplementary Table 1

*Details of the sampling sites including site code, geographic locations [West (W), South (S) and East (E)], sampling date (year 2017) and type of site (natural or artificial). Order of appearance of sites progresses along the sampled coast, from the northwest coast to the northeast coast of South Africa.*

| Site Name | Site Code | Latitude (S) | Longitude (E) | Sampling Date | Geographic Coastline | Site Classification |
| --- | --- | --- | --- | --- | --- | --- |
| **Yachtport SA, Saldanha** | SY | -33.026701 | 17.960758 | 14-Oct | W | Artificial |
| **Saldanha Natural Site** | SN | -33.071672 | 18.038446 | 15-Oct | W | Natural |
| **Club Mykonos Marina** | SM | -33.046315 | 18.04028 | 15-Oct | W | Artificial |
| **V&A Waterfront Marina** | TB | -33.909055 | 18.419893 | 18-Oct | W | Artificial |
| **Hout Bay Yacht Club** | HB | -34.049767 | 18.348042 | 19-Oct | W | Artificial |
| **Hout Bay Natural Site** | HN | -34.049016 | 18.360954 | 19-Oct | W | Natural |
| **Mossel Bay Marina** | MB | -34.178386 | 22.144918 | 22-Oct | S | Artificial |
| **Mossel Bay Natural Site** | MN | -34.299278 | 21.944026 | 22-Oct | S | Natural |
| **Knysna** | KN | -34.041086 | 23.043592 | 24-Oct | S | Artificial |
| **Knysna Natural Site** | NK | -34.05877 | 23.034025 | 24-Oct | S | Natural |
| **Cape St Francis** | CN | -34.212257 | 24.837625 | 26-Oct | S | Natural |
| **Port Elizabeth Marina** | PE | -33.96692 | 25.634461 | 27-Oct | S | Artificial |
| **Bushmans River (Kenton Marina)** | BR | -33.679558 | 26.655841 | 30-Oct | E | Artificial |
| **Port Alfred Marina** | PA | -33.593546 | 26.892084 | 31-Oct | E | Artificial |
| **East London Marina** | EL | -33.024163 | 27.896269 | 02-Nov | E | Artificial |
| **Durban Marina** | DU | -29.862663 | 31.021903 | 08-Nov | E | Artificial |
| **Richards Bay Marina** | RB | -28.793908 | 32.079184 | 10-Nov | E | Artificial |
| **Richards Bay Natural** | RN | -28.827437 | 32.035825 | 11-Nov | E | Natural |

### Supplementary Table 2

*Descriptive statistics of bioinformatic parameters from environmental DNA metabarcoding data of natural and artificial sites along the coast of South Africa. Each column indicates* **A** *the gene fragment used in the metabarcoding experiment; COI – cytochrome c oxidase subunit I; 18S -nuclear small subunit ribosomal DNA, 16S – prokaryotic small subunit ribosomal DNA* **B** *the dataset subset into metazoan (COI), protist (18S) and bacterial (16S) datasets. The first two rows indicate the number of reads and standard deviation per sample. The third indicates the lowest number of reads per sample used to rarefy the data for analyses. The fourth and fifth rows indicate the number of Amplicon Sequence Variants (ASVs) from the dataset and the number that did not have a (70% percentage identity) match in the NCBI nt database. The final row indicates the number of ASVs with high quality assignment, the values in brackets identify the percentage identity used for high quality assignment.*

**A**

|  | **COI** | **18S** | **16S** |
| --- | --- | --- | --- |
| **Raw mean reads per sample** | 207,675.8 | 361,868.6 | 473,062.8 |
| **Standard Deviation** | 112,122.6 | 110,399.3 | 106,473.1 |
| **N_reads_ Rarefication** | 61,958 | 147,307 | 115,960 |
| **ASVs in analysis** | 4,867 | 2,364 | 2,826 |
| **ASVs with no taxonomic assignment** | 2,824 | 131 | 8 |
| **ASVs with high quality BLAST nt assignment** | 150 *(97% ID)* | 265 *(99% ID)* | 1,660 *(Genus assignment)* |

**B**

|  | **Metazoa** | **Protista** | **Bacteria** |
| --- | --- | --- | --- |
| **N_reads_ Rarefication** | 8,682 | 12,161 | 115,959 |
| **ASVs in analysis** | 1,054 | 1,433 | 2,826 |

Supplementary Table 3

*Analysis of variance (ANOVA) testing differences in amplicon sequence variant richness between coasts for datasets by* ***A*** *taxa and* ***B*** *marker, a Tukey post-hoc test for 16S is shown.*

*Significant p-values at α=0.05 level are indicated in* ***bold****.*

**A**

| **ANOVA** | Df | SumofSquares | MeanSquare | F | P value |
| --- | --- | --- | --- | --- | --- |
| **Metazoa** |  |  |  |  |  |
| Coast | 2 | 4054 | 2027 | 1.941 | 0.178 |
| Residuals | 15 | 15664 | 1044 |  |  |
| **Protista** |  |  |  |  |  |
| Coast | 2 | 7819 | 3910 | 1.416 | 0.273 |
| Residuals | 15 | 41404 | 2760 |  |  |
| **Bacteria** |  |  |  |  |  |
| Coast | 2 | 79844 | 39922 | 7.178 | **0.007** |
| Residuals | 15 | 83422 | 5561 |  |  |
| **TukeyPostHoc** | |  |  |  |  |
|  | **Bacteria** |  |  |  |  |
|  |  | diff | lwr | upr | P value |
|  | South-East | 146.167 | 34.330 | 258.004 | **0.012** |
|  | West-East | 10.333 | -101.504 | 122.170 | 0.969 |
|  | West-South | -135.833 | -247.670 | -23.996 | **0.017** |

**B**

| **ANOVA** | Df | SumofSquares | MeanSquare | F | P value |
| --- | --- | --- | --- | --- | --- |
| **COI** |  |  |  |  |  |
| Coast | 2 | 0.115 | 0.05763 | 3.64 | 0.051 |
| Residuals | 15 | 0.237 | 0.01583 |  |  |
| **18S** |  |  |  |  |  |
| Coast | 2 | 28434 | 14217 | 1.824 | 0.195 |
| Residuals | 15 | 116945 | 7796 |  |  |
| **16S** |  |  |  |  |  |
| Coast | 2 | 79844 | 39922 | 7.178 | **0.007** |
| Residuals | 15 | 83422 | 5561 |  |  |

Supplementary Table 4

*Permutational analysis of variance (PERMANOVA) model outputs based on a Jaccard index of amplicon sequence variants from eDNA metabarcoding of sites across South Africa. Models are presented for* **A** *three taxonomic groups; metazoa, protista and bacteria.* **B** *Three genes; COI – cytochrome c oxidase subunit I; 18S -nuclear small subunit ribosomal DNA, 16S – prokaryotic small subunit ribosomal DNA.*

*Significant p-values at α=0.05 level are indicated in* ***bold****.*

**A**

|  | Df | SumsOfSqs | MeanSqs | Pseudo F | R^2^ | P value |
| --- | --- | --- | --- | --- | --- | --- |
| **Metazoa** |  |  |  |  |  |  |
| Coast | 2 | 1.511 | 0.755 | 1.981 | 0.209 | **0.001** |
| Residuals | 15 | 5.721 | 0.381 |  | 0.791 |  |
| Total | 17 | 7.232 |  |  |  |  |
| **Protists** |  |  |  |  |  |  |
| Coast | 2 | 1.601 | 0.800 | 2.656 | 0.262 | **0.001** |
| Residuals | 15 | 4.520 | 0.302 |  | 0.738 |  |
| Total | 17 | 6.121 |  |  |  |  |
| **Bacteria** |  |  |  |  |  |  |
| Coast | 2 | 1.737 | 0.867 | 3.229 | 0.301 | **0.001** |
| Residuals | 15 | 4.035 | 0.269 |  | 0.699 |  |
| Total | 17 | 5.773 |  |  |  |  |

**B**

|  | Df | SumsOfSqs | MeanSqs | Pseudo F | R^2^ | P value |
| --- | --- | --- | --- | --- | --- | --- |
| **COI** |  |  |  |  |  |  |
| Coast | 2 | 1.632 | 0.816 | 2.083 | 0.217 | **0.001** |
| Residuals | 15 | 5.879 | 0.392 |  | 0.784 |  |
| Total | 17 | 7.510 |  |  |  |  |
| **18S** |  |  |  |  |  |  |
| Coast | 2 | 1.192 | 0.596 | 1.358 | 0.153 | **0.001** |
| Residuals | 15 | 6.580 | 0.439 |  | 0.845 |  |
| Total | 17 | 7.772 |  |  |  |  |
| **16S** |  |  |  |  |  |  |
| Coast | 2 | 1.670 | 0.835 | 2.712 | 0.267 | **0.001** |
| Residuals | 15 | 4.618 | 0.308 |  | 0.734 |  |
| Total | 17 | 6.287 |  |  |  |  |

Supplementary Table 5

*Pairwise permutational analysis of variance (PERMANOVA) model outputs based on a Jaccard index of amplicon sequence variants from eDNA metabarcoding of sites across South Africa. Models are presented for three genes; COI - cytochrome c oxidase subunit I; 18S - nuclear small subunit ribosomal DNA, 16S - prokaryotic small subunit ribosomal DNA. Significant p-values at α=0.05 level are indicated in* ***bold****.*

**A**

|  | Df | SumsOfSqs | F.Model | R2 | p.value | p.adjusted |
| --- | --- | --- | --- | --- | --- | --- |
| **Metazoa** |  |  |  |  |  |  |
| East vs South | 1 | 0.671 | 1.853 | 0.156 | 0.002 | **0.005** |
| East vs West | 1 | 0.859 | 2.222 | 0.182 | 0.005 | **0.005** |
| South vs West | 1 | 0.737 | 1.861 | 0.157 | 0.005 | **0.005** |
| **Protista** |  |  |  |  |  |  |
| East vs South | 1 | 0.800 | 2.843 | 0.221 | 0.004 | **0.004** |
| East vs West | 1 | 0.885 | 2.887 | 0.224 | 0.004 | **0.004** |
| South vs West | 1 | 0.717 | 2.266 | 0.185 | 0.003 | **0.004** |
| **Bacteria** |  |  |  |  |  |  |
| East vs South | 1 | 0.738 | 3.135 | 0.239 | 0.001 | **0.003** |
| East vs West | 1 | 1.093 | 3.993 | 0.285 | 0.003 | **0.003** |
| South vs West | 1 | 0.775 | 2.602 | 0.206 | 0.003 | **0.003** |

**B**

|  | Df | SumsOfSqs | F.Model | R2 | p.value | p.adjusted |
| --- | --- | --- | --- | --- | --- | --- |
| **COI** |  |  |  |  |  |  |
| East vs South | 1 | 0.705 | 1.723 | 0.147 | 0.002 | **0.006** |
| East vs West | 1 | 0.899 | 2.232 | 0.182 | 0.003 | **0.009** |
| South vs West | 1 | 0.844 | 2.322 | 0.188 | 0.001 | **0.003** |
| **18S** |  |  |  |  |  |  |
| East vs South | 1 | 0.612 | 1.411 | 0.124 | 0.016 | **0.048** |
| East vs West | 1 | 0.615 | 1.374 | 0.121 | 0.002 | **0.006** |
| South vs West | 1 | 0.560 | 1.290 | 0.114 | 0.006 | **0.018** |
| **16S** |  |  |  |  |  |  |
| East vs South | 1 | 0.732 | 2.180 | 0.179 | 0.007 | **0.021** |
| East vs West | 1 | 1.081 | 3.411 | 0.254 | 0.004 | **0.012** |
| South vs West | 1 | 0.691 | 2.553 | 0.203 | 0.002 | **0.006** |

### Supplementary Table 6

*Analysis of variance model output for tests of multivariate homogeneity of group dispersion (betadisp R function) based on a Jaccard index of amplicon sequence variant s from eDNA metabarcoding of sites across South Africa. Models are presented for three genes; COI – cytochrome c oxidase subunit I; 18S -nuclear small subunit ribosomal DNA, 16S – prokaryotic small subunit ribosomal DNA.*

*Significant p-values at α=0.05 level are indicated in* ***bold****.*

**A**

|  | Df | SumsOfSqs | MeanSq | F.Value | P |
| --- | --- | --- | --- | --- | --- |
| **Metazoa** |  |  |  |  |  |
| Groups | 2 | 0.008469 | 0.0042347 | 1.9485 | 0.177 |
| Residuals | 15 | 0.032599 | 0.0021733 |  |  |
| **Protista** |  |  |  |  |  |
| Groups | 2 | 0.012200 | 0.0061001 | 1.4672 | 0.262 |
| Residuals | 15 | 0.062364 | 0.0041576 |  |  |
| **Bacteria** |  |  |  |  |  |
| Groups | 2 | 0.038883 | 0.0194414 | 4.0874 | **0.038** |
| Residuals | 15 | 0.071346 | 0.0047564 |  |  |
| **TukeyPostHoc**  **(Bacteria/16S)** | Comparison | diff | lwr | upr | P.adj |
|  | South vs West | 0.04778014 | -0.0556455 | 0.1512058 | 0.471 |
|  | East vs West | 0.11338036 | 0.00995475 | 0.216806 | **0.031** |
|  | East vs South | 0.06560021 | -0.0378254 | 0.1690258 | 0.257 |

**B**

|  | Df | SumsOfSqs | MeanSq | F.Value | P |
| --- | --- | --- | --- | --- | --- |
| **COI** |  |  |  |  |  |
| Groups | 2 | 0.016455 | 0.0082276 | 2.3376 | 0.1307 |
| Residuals | 15 | 0.052731 | 0.0035154 |  |  |
| **18S** |  |  |  |  |  |
| Groups | 2 | 0.0014773 | 0.00073863 | 0.5964 | 0.5633 |
| Residuals | 15 | 0.0185756 | 0.00123837 |  |  |
| **16S** |  |  |  |  |  |
| Groups | 2 | 0.037211 | 0.0186056 | 3.9383 | **0.0422** |
| Residuals | 15 | 0.070864 | 0.0047243 |  |  |

### Supplementary Table 7

*Mantel test summary output for both Partial mantel and corrected Mantel tests. Parameters are as follows, SST – mean sea surface temperature (°C); SSS – mean sea surface salinity (parts per thousand); Chl a – chlorophyll a concentration (mg m^-3^); impact – human marine impact score (unitless measurement, see details in main text). Models are presented for three genes; COI – cytochrome c oxidase subunit I; 18S -nuclear small subunit ribosomal DNA, 16S – prokaryotic small subunit ribosomal DNA.*

*Significant p-values at α=0.05 level are indicated in* ***bold****.*

**A**

|  | **SST** |  | **SSS** |  | **ChlA** |  | **Impact** |  |
| --- | --- | --- | --- | --- | --- | --- | --- | --- |
|  | **R statistic** | **p value** | **R statistic** | **p value** | **R statistic** | **p value** | **R statistic** | **p value** |
| **Metazoans partial** | 0.264 | **0.013** | -0.148 | 0.881 | 0.066 | 0.272 | 0.432 | **0.001** |
| **Metazoans corrected** | 0.175 | **0.015** | -0.031 | 0.65 | 0.071 | 0.235 | 0.308 | **0.002** |
| **Protists partial** | 0.135 | 0.073 | 0.026 | 0.388 | 0.214 | **0.012** | 0.317 | **0.001** |
| **Protists corrected** | 0.129 | 0.066 | 0.04 | 0.352 | 0.165 | **0.04** | 0.249 | **0.01** |
| **Bacteria partial** | 0.304 | **0.008** | -0.133 | 0.843 | 0.021 | 0.403 | 0.492 | **0.001** |
| **Bacteria corrected** | 0.207 | **0.007** | -0.001 | 0.536 | 0.062 | 0.294 | 0.326 | **0.002** |

**B**

|  | **SST** |  | **SSS** |  | **ChlA** |  | **Impact** |  |
| --- | --- | --- | --- | --- | --- | --- | --- | --- |
|  | **R statistic** | **p value** | **R statistic** | **p value** | **R statistic** | **p value** | **R statistic** | **p value** |
| **Partial Mantel COI** | 0.323 | **0.002** | 0.002 | 0.484 | 0.077 | 0.195 | 0.357 | **0.003** |
| **Corrected Mantel COI** | 0.157 | **0.022** | 0.005 | 0.531 | 0.057 | 0.277 | 0.234 | **0.007** |
| **Partial Mantel 18S** | 0.143 | **0.059** | -0.024 | 0.602 | 0.139 | 0.076 | 0.394 | **0.001** |
| **Corrected Mantel 18S** | 0.162 | **0.03** | 0.041 | 0.355 | 0.134 | 0.087 | 0.297 | **0.003** |
| **Partial Mantel 16S** | 0.304 | **0.008** | -0.133 | 0.842 | 0.021 | 0.407 | 0.492 | **0.001** |
| **Corrected Mantel 16S** | 0.207 | **0.007** | 0 | 0.531 | 0.062 | 0.302 | 0.326 | **0.002** |

### Supplementary Table 8

*Distance based redundancy analysis model outputs for models with Jaccard dissimilarities from eDNA metabarcoding data from South Africa as the response variable. Explanatory variables are as follows, SST – mean sea surface temperature (°C); Chl a – chlorophyll a concentration (mg m^-3^); impact – human marine impact score (unitless measurement, see details in main text). Models are presented for datasets composed of metazoa, protista and bacteria.*

*Significant p-values at α=0.05 level are indicated in* ***bold****.*

|  | Df | SumOfSqs. | F Value | P Value |
| --- | --- | --- | --- | --- |
| **Metazoa** |  |  |  |  |
| SST | 1 | 0.8328 | 2.1424 | **1.00E-04** |
| Impact | 1 | 0.5165 | 1.3286 | **0.0277** |
| Residual | 15 | 5.8308 |  |  |
|  |  | Adj.R2 | 0.086 |  |
| **Protista** |  |  |  |  |
| Chl a | 1 | 0.7888 | 2.4641 | **1.00E-04** |
| Impact | 1 | 0.5301 | 1.656 | **0.009299** |
| Residual | 15 | 4.802 |  |  |
|  |  | Adj.R2 | 0.111 |  |
| **Bacteria** |  |  |  |  |
|  | Df | SumOfSqs | F | Pr(>F) |
| SST | 1 | 1.0178 | 3.655 | **1.00E-04** |
| Impact | 1 | 0.5194 | 1.8652 | **0.009299** |
| Residual | 15 | 4.1769 |  |  |
|  |  | Adj.R2 | 0.18 |  |
| **Metazoa (18S)** | |  |  |  |
|  | Df | SumOfSqs | F | Pr(>F) |
| SST | 1 | 0.823 | 2.2557 | **1.00E-04** |
| Impact | 1 | 0.6059 | 1.6607 | **0.0021** |
| Residual | 15 | 5.4728 |  |  |
|  |  | Adj.R2 | 0.111 |  |
| **Protista (COI)** | |  |  |  |
|  | Df | SumOfSqs | F | Pr(>F) |
| SST | 1 | 1.2049 | 3.9839 | **1.00E-04** |
| Impact | 1 | 0.5637 | 1.8638 | **0.0171** |
| Residual | 15 | 4.5366 |  |  |
|  |  | Adj.R2 | 0.192 |  |

### Supplementary Table 9

*Generalised additive model with a restricted maximum likelihood (REML) 2D smoother summary, each row represents an individually fitted model. Parameters are as follows, SST – mean sea surface temperature (°C); SSS – mean sea surface salinity (parts per thousand); Chl a – chlorophyll a concentration (mg m^-3^); impact – human marine impact score (unitless measurement, see details in main text). Models are presented for three genes; COI – cytochrome c oxidase subunit I; 18S -nuclear small subunit ribosomal DNA, 16S – prokaryotic small subunit ribosomal DNA.*

*Significant p-values at α=0.05 level are indicated in* ***bold****.*

**A**

|  | Parametric Coef. | |  |  | Smoother | |  |  |
| --- | --- | --- | --- | --- | --- | --- | --- | --- |
|  | Estimate | Std.Error | T Value | P Value | Df | F | P Value | Adj.R2 |
| Metazoa |  |  |  |  |  |  |  |  |
| Impact | 3.2348 | 0.1593 | 20.31 | 2.08E-10 | 9 | 3.563 | **0.00297** | 0.654 |
| SST | 18.56054 | 0.09681 | 191.7 | <2e-16 | 9 | 100.2 | **<2e-16** | 0.982 |
| Protista |  |  |  |  |  |  |  |  |
| impact | 3.2348 | 0.1779 | 18.18 | 5.8E-10 | 9 | 2.481 | **0.0126** | 0.568 |
| chl a | 1.8003 | 0.05603 | 32.13 | 1.89E-14 | 9 | 16.15 | 2.63E-13 | 0.895 |
| Bacteria |  |  |  |  |  |  |  |  |
| impact | 3.2348 | 0.1482 | 21.83 | 2.45E-10 | 9 | 4.409 | **0.00207** | 0.700 |
| SST | 18.56054 | 0.09753 | 190.3 | <2e-16 | 9 | 98.73 | **<2e-16** | 0.981 |

**B**

|  | Parametric Coef. | |  |  | Smoother | |  |  |
| --- | --- | --- | --- | --- | --- | --- | --- | --- |
|  | Estimate | Std.Error | T Value | P Value | Df | F | P Value | Adj.R2 |
| COI |  |  |  |  |  |  |  |  |
| SSS | 35.225667 | 0.006731 | 5233 | <2e-16 | 9 | 27.48 | **3.79E-13** | 0.936 |
| impact | 3.2348 | 0.1753 | 18.45 | 1.51E-10 | 9 | 2.608 | **0.0044** | 0.58 |
| chl a | 1.8003 | 0.05529 | 32.56 | 3.04E-13 | 9 | 16.63 | **3.47E-12** | 0.898 |
| SST | 18.56054 | 0.09966 | 186.2 | <2e-16 | 9 | 94.47 | **<2e-16** | 0.98 |
| 18S |  |  |  |  |  |  |  |  |
| SSS | 35.225667 | 0.008706 | 4046 | <2e-16 | 9 | 15.67 | **5.68E-09** | 0.892 |
| impact | 3.2348 | 0.1477 | 21.9 | 9.75E-11 | 9 | 4.45 | **0.00117** | 0.702 |
| chl a | 1.8003 | 0.04337 | 41.51 | 3.86E-16 | 9 | 28.21 | **<2e-16** | 0.937 |
| SST | 18.5605 | 0.1531 | 121.2 | <2e-16 | 9 | 38.93 | **<2e-16** | 0.954 |
| 16S |  |  |  |  |  |  |  |  |
| SSS | 35.225667 | 0.007869 | 4476 | <2e-16 | 9 | 19.6 | **1.00E-10** | 0.912 |
| impact | 3.2348 | 0.1545 | 20.94 | 2.93E-10 | 9 | 3.907 | **0.00311** | 0.674 |
| chl a | 1.8003 | 0.07156 | 25.16 | 3.76E-12 | 9 | 9.17 | **1.47E-07** | 0.829 |
| SST | 18.56054 | 0.09916 | 187.2 | <2e-16 | 9 | 95.44 | **<2e-16** | 0.981 |

### Supplementary Figure 1

*Bar charts indicating the proportion of reads assigned per phyla(metazoans/bacteria) or supergroup (protists) from environmental DNA metabarcoding of seawater collected from sites across South Africa. The three rows correspond with data from metazoans (top), protists (middle) and bacteria (bottom). Site name abbreviations as in Supplementary Table 1.*

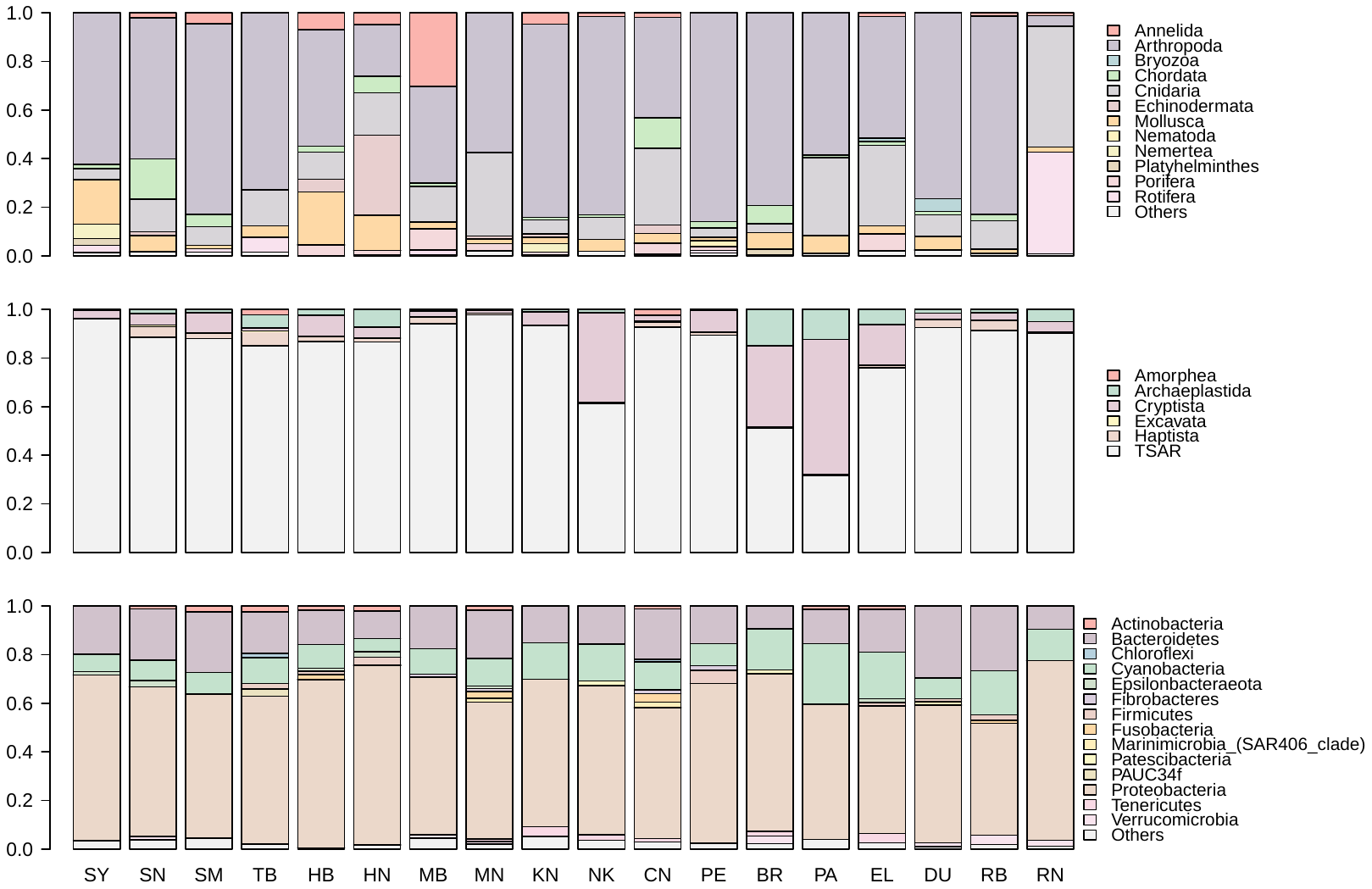

### Supplementary Figure 2

*Observed patterns of β-diversity from environmental DNA metabarcoding of:* ***a*** *metazoans from the 18S dataset and* ***b*** *protists from the COI dataset; based on Jaccard dissimilarities between amplicon sequence variants along the coast of South Africa. The first column of plots shows non-metric multidimensional scaling (nMDS) ordinations. Coloured hulls show the spread of the data and lines indicate the spread around the centroid grouped by coast with the east, south and west coasts denoted by orange, green and blue respectively. Site name abbreviations as in Supplementary Table 1. Natural sites are denoted with triangles and artificial sites with filled circles. The second column of plots shows the same nMDS ordinations as the first column including the output of a generalised additive model with a 2D smoothed function for each of the significant environmental / impact variables overlaid; temperature – mean sea surface temperature (°C); Chlorophyll a – chlorophyll a concentration (mg m^-3^); impact – human marine impact score (unitless measurement, see details in main text) against the two nMDS axes. The venn diagram charts indicate the percentage total of variance in community dissimilarity explained by each significant variable, derived using variance partitioning of a distance-based redundancy analysis.*

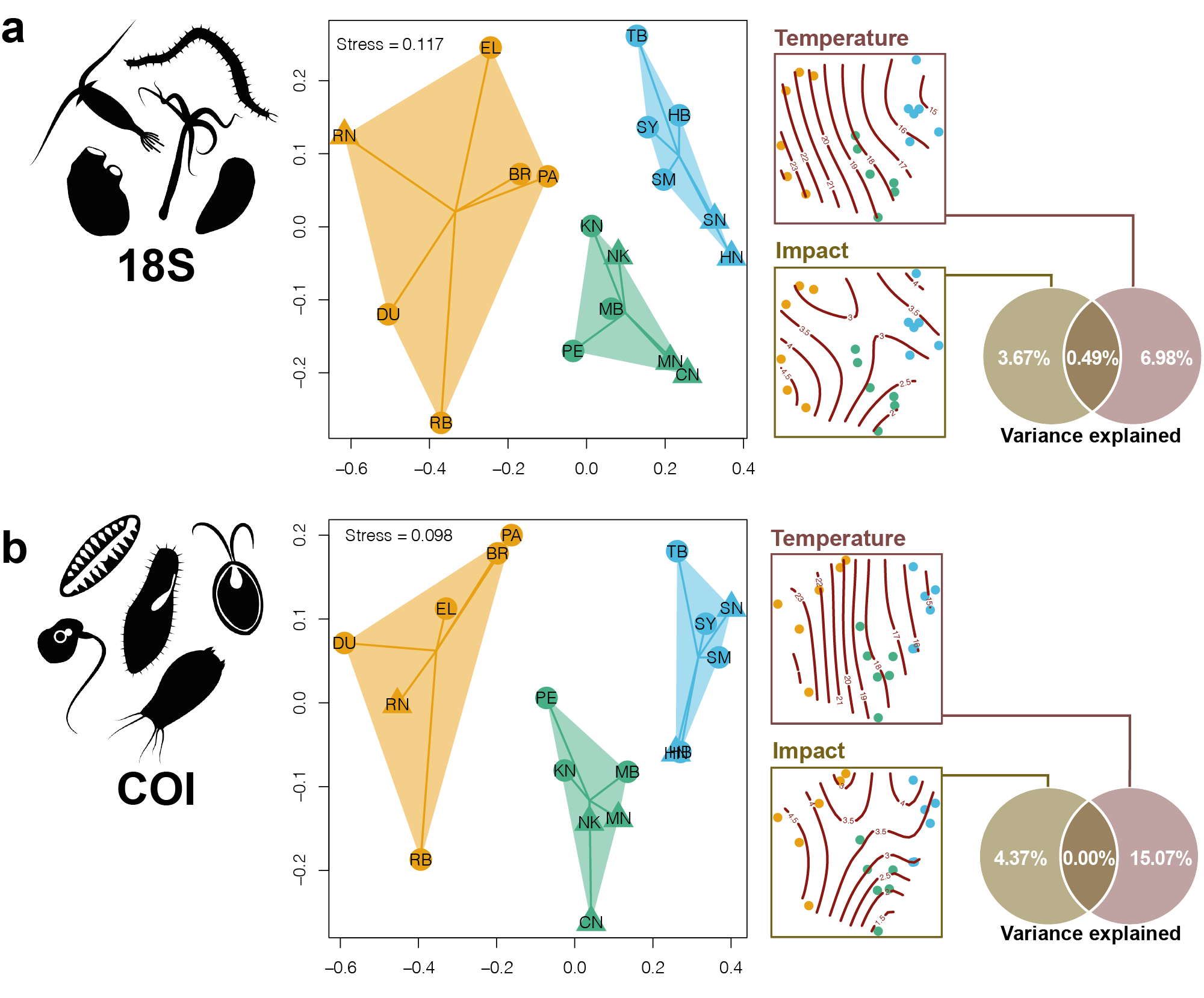

### Supplementary Information 1

*Quality control method and results for detecting non-biological (chimeras, unknown primer dimers, etc.) and biological (numts, pseudogenes, non-region binding) erroneous amplicon sequence variants (ASVs) in metabarcoding of eDNA data from South African seawater.*

**Open reading frame trial**

Each ASV in the raw (post DADA2 pipeline) and quality controlled (post LULU and read frequency filtering) COI dataset was translated at each of three reading frames under all possible translation tables. The smallest number of stop codons for each frame/table was recorded per ASV. In addition, the number of stop codons per ASV for the invertebrate mitochondrial codon table was recorded to evaluate the possibility that using the minimum value across all tables ignores true stop codons as a result of using a table with fewer stop codons (e.g. NCBI Table 14 – the alternative flatworm table). The number of stop codons per ASV and percentage of ASVs with no stop codon is shown in the table below.

|  | QC Dataset | | Raw Dataset | |
| --- | --- | --- | --- | --- |
| Nstop Codons | MinStop | Invert | MinStop | Invert |
| 0 | 4866 | 4845 | 37242 | 36923 |
| 1 | 1 | 11 | 72 | 138 |
| 2 | 0 | 5 | 2 | 100 |
| 3 | 0 | 5 | 0 | 85 |
| 4 | 0 | 0 | 0 | 54 |
| 5 | 0 | 1 | 0 | 15 |
| 6 | 0 | 0 | 0 | 0 |
| 7 | 0 | 0 | 0 | 1 |
| Percentage of stop | 99.98 | 99.55 | 99.80 | 98.95 |

Overall, the proportion of sequences in the QC dataset with a stop codon was negligible, indicating that only a minute fraction of the observed sequences are non-coding.

**Manual BLAST trial**

A random 5% subset of ASVs (243 sequences) in the quality-controlled cytochrome c oxidase I (COI) dataset was extracted. These ASVs were compared to the NCBI nt/nr database using the megablast algorithm on the online NCBI blast tool (blast.ncbi.nlm.nih.gov, accessed online 07.04.2020). Each sequence was manually examined, the top 50 hits were visually compared, and the best match (highest E value above 75% coverage) was recorded. Any ASV with a less than 70% coverage by a subject sequence was treated as a putative chimera. The query section not matching the subject in these cases was extracted and used as a query in a second search. A true chimera would have different biological hits for each section. For ASVs where no hit was found in the above database, a blastx search (nucleotide -> protein search) was performed under the standard genetic code. Protein sequence matches were expected to be cytochrome c oxidase I (COI), hits with less than 97% positive ID (amino acid matches that are identical or have conservative substitutions) were designated a ‘low’ quality match.

Out of 243 sequences, 20 (8.2%) could be assigned a high-quality hit of 97% sequence identity or greater. 216 (88.8%) could be assigned a poor-quality match indicating a species or taxa not represented in the sequence database. 2 (0.8%) sequences had no nucleotide match but a high-quality protein match indicating a functioning COI gene. Five sequences (2.0%) had no nucleotide match and no high-quality protein match, however all five sequences received protein matches to a COI subject indicating biological rather than technical origin. No sequences had a partial (<70%) match indicative of a chimeric sequence. Overall, this evidence suggests that the bioinformatic parameters and filtering steps used to produce the quality-controlled dataset resulted in a tiny minority of retained sequences that correspond with technical errors and non-target biological regions.

### Supplementary Information 2

*Information concerning control samples used during field sampling and in the laboratory during the construction of metabarcoding libraries for high-throughput sequencing of eDNA extracted from water samples collected along the coast of South Africa.*

Each water sample was taken using a peristaltic pump, which was washed with 1 litre of seawater from the site before conducting the field sampling. While in the laboratory, all reused field equipment (hosing and plasticware) was washed thoroughly with 5% bleach solution and rinsed with tap water.

Negative controls were implemented during each DNA extraction and during each PCR step of library construction, with DNA extractions and PCRs processed with PCR grade water instead of filters or water. Across all controls for COI a total of 6.6 ± 7.6 (s.d.) ASVs were found in each control sample with a summed average of 117.3 ± 155.1 reads per sample. Across the 18S controls 12.1 ± 9.5 (s.d.) ASVs with a summed average of 579.1 ± 517.5 reads per sample. The 16S data showed the largest number of average ASVs (61.7 ± 63.1) per sample and also more summed reads on average (16,079.9 ± 31,915.2). A single ASV, assigned to the genus *Bradyrhizobium*, contributed the bulk of these reads (mean of 25,444.4 per control sample) and was found only in control samples used to assess decontamination in field. This genus of bacteria are Gram-negative soil bacteria, some of which have roles in N_2_ fixation (Stacey et al. 1995) and therefore likely represent contamination from the water used for cleaning apparatus between sites. However, this genus of bacteria has also been shown to contaminate ultra-pure water and laboratory reagents (Kulakov et al. 2002, Salter et al. 2014). After bioinformatically removing this ASV the 16S control samples contained a mean of 56.1 ± 61.9 ASVs per sample and a mean of 3,054.9 ± 3,398.8 summed reads per sample.

The proportion of contamination assigned to travel, lab and PCR controls are shown below in Fig. SI.21. The largest proportion of contamination was from the travel controls, indicating that the cleaning process between sites was the largest source of contamination. The level of contamination shown here is typical for eDNA studies (Holman et al. 2019, Jeunen et al. 2019, Blackman et al. 2020) and, as all contamination was bioinformatically filtered as detailed in the main manuscript. Therefore, we are confident that such contamination did not affect the inferences presented in the main manuscript.

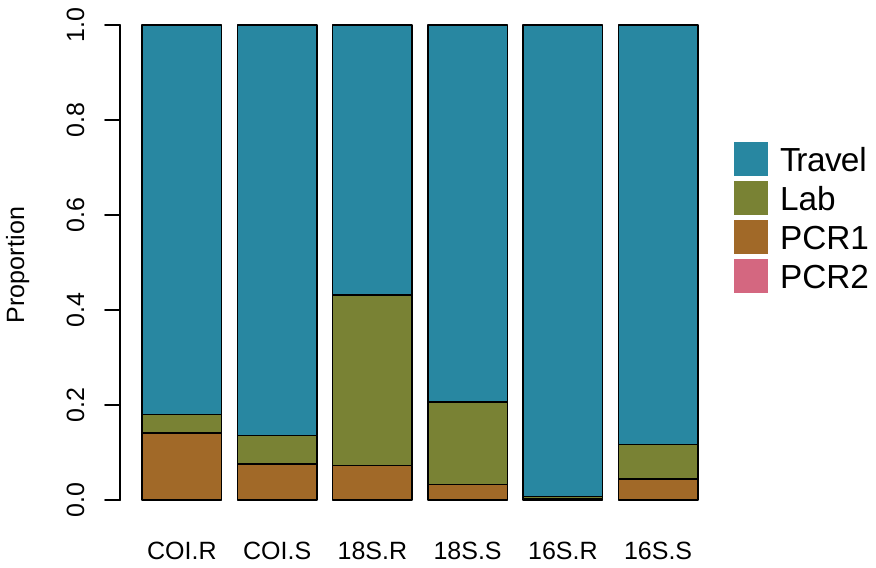

***Figure SI.21*** *Proportion of contamination attributed to different sections of the workflow. Each column shows the proportion of either reads (COI.R, 18.R, 16.R) or ASVs (COI.S, 18S.S, 16S.S) contributed by each of different types of control sample for all three sequenced markers (COI, 18S, 16S). Travel controls include sealed controls and tap water controls used to evaluate cleaning of equipment between sampling. Lab controls include sealed filters, DNA extraction kits and inhibition clean up kits. PCR controls are no template controls added before PCR.*

### Supplementary Information 3

*Phyla-level analysis of biogeographic patterns*

In order to evaluate the beta diversity patterns at lower taxonomic levels than kingdom each dataset was subset to include only ASVs allocated to the top five most ASV rich phyla per taxonomic dataset (metazoans, protists, bacteria). A non-metric multidimensional ordination, evaluation of multivariate heterogeneity and PERMANOVA were performed on each of these sub-datasets as detailed in the main manuscript methods for the complete datasets.

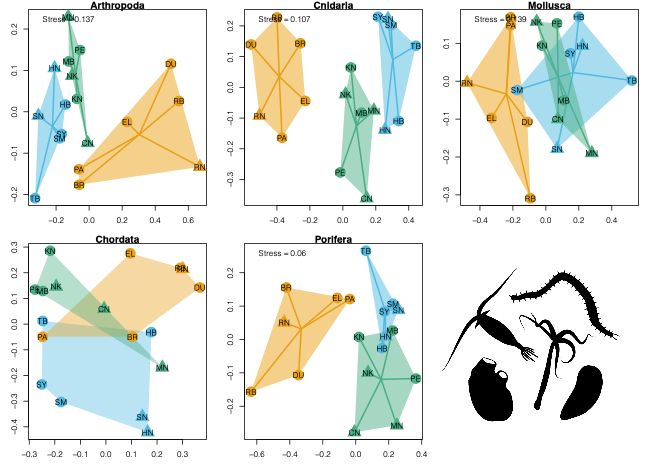

***Figure SI.31*** *Observed patterns of β-diversity from environmental DNA metabarcoding (COI) based on Jaccard dissimilarities between amplicon sequence variants along the coast of South Africa. Plots shows non-metric multidimensional scaling (nMDS) ordinations for each phylum. Coloured hulls show the spread of the data and lines indicate the spread around the centroid grouped by coast with the east, south and west coasts denoted by orange, green and blue respectively. Site name abbreviations as in Supplementary Table 1. The ordination for Chordata is the first two axes of a principle coordinate analysis due to problems with convergence of an nMDS solution that well represents the Chordata data.*

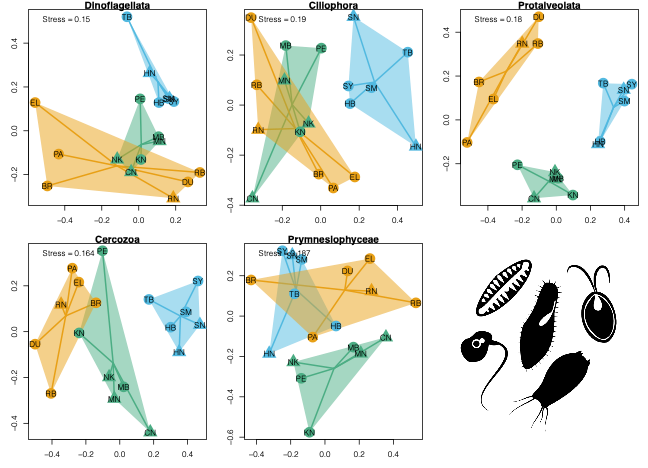

***Figure SI.32*** *Observed patterns of β-diversity from environmental DNA metabarcoding (18S) based on Jaccard dissimilarities between amplicon sequence variants along the coast of South Africa. Plots shows non-metric multidimensional scaling (nMDS) ordinations for each phylum. Coloured hulls show the spread of the data and lines indicate the spread around the centroid grouped by coast with the east, south and west coasts denoted by orange, green and blue respectively. Site name abbreviations as in Supplementary Table 1.*

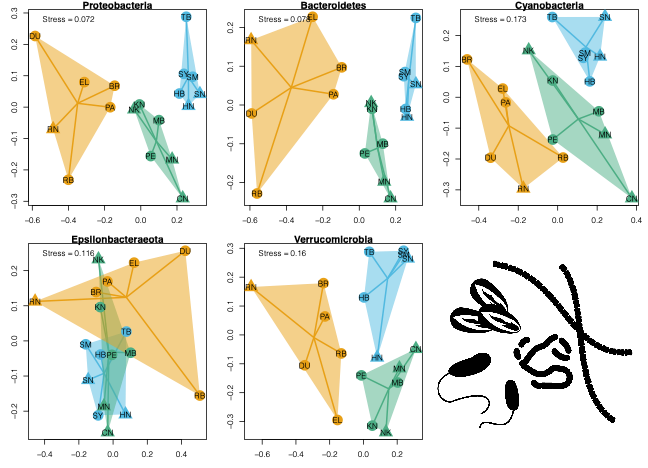

***Figure SI.31*** *Observed patterns of β-diversity from environmental DNA metabarcoding (16S) based on Jaccard dissimilarities between amplicon sequence variants along the coast of South Africa. Plots shows non-metric multidimensional scaling (nMDS) ordinations for each phylum. Coloured hulls show the spread of the data and lines indicate the spread around the centroid grouped by coast with the east, south and west coasts denoted by orange, green and blue respectively. Site name abbreviations as in Supplementary Table 1.*

***Table SI.31*** *Phyla level dataset outputs showing number of ASVs per phyla (nASVs), the outputs (F statistic and p value) from an analysis of variance on multivariate distribution between ecoregions for each dataset (permdisp from vegan in R). F statistic and P value from a PERMANOVA model based on Jaccard dissimilarities testing a difference in between ecoregions.*

*Significant p-values at α=0.05 level are indicated in* ***bold****.*

|  |  |  | **Multivariate Dispersion** | | **PERMANOVA** | |
| --- | --- | --- | --- | --- | --- | --- |
| **Phyla** | **Dataset** | **nASVs** | **Fstat** | **p value** | **Fstat** | **p value** |
| **Arthropoda** | Metazoa | 354 | 1.742 | 0.209 | 1.816 | **0.001** |
| **Cnidaria** | Metazoa | 258 | 0.774 | 0.479 | 2.468 | **0.001** |
| **Mollusca** | Metazoa | 106 | 0.377 | 0.692 | 2.07 | **0.001** |
| **Chordata** | Metazoa | 92 | 0.241 | 0.789 | 1.452 | **0.001** |
| **Porifera** | Metazoa | 68 | 2.771 | 0.095 | 2.261 | **0.001** |
| **Dinoflagellata** | Protist | 257 | 2.637 | 0.104 | 3.086 | **0.001** |
| **Ciliophora** | Protist | 389 | 0.437 | 0.654 | 2.024 | **0.001** |
| **Protalveolata** | Protist | 281 | 1.958 | 0.176 | 2.802 | **0.001** |
| **Cercozoa** | Protist | 218 | 1 | 0.391 | 2.446 | **0.001** |
| **Prymnesiophyceae** | Protist | 70 | 0.178 | 0.839 | 2.598 | **0.001** |
| **Proteobacteria** | Bacteria | 1340 | 2.489 | 0.117 | 3.208 | **0.001** |
| **Bacteroidetes** | Bacteria | 673 | 7.38 | **0.006** | 4.084 | **0.001** |
| **Cyanobacteria** | Bacteria | 390 | 2.776 | 0.094 | 2.826 | **0.001** |
| **Epsilonbacteraeota** | Bacteria | 83 | 2.158 | 0.15 | 1.47 | **0.029** |
| **Verrucomicrobia** | Bacteria | 84 | 0.45 | 0.646 | 2.748 | **0.001** |

### Supplementary Information 4

*Simulations of power for detecting differences in eco-regions using PERMANOVAs.*

We tested how many species observations were required to detect a significant difference between the multivariate spread of the three ecoregions (see Methods, main text). This was important as the number of ASVs subsets at phyla level can be low (some phyla had less than 20 observations, see Table SI.31). Therefore, we conducted a community simulation using data simulated using random distributions in R (v4.0.2) as follows.
A randomly-generated dataset containing 1000 simulated species was created. Three possible types of spatial distribution were possible for each species: 1. Panmixia (i.e. the species is present in all sites), 2. Random distribution (randomly distributed across sites), and 3. ecoregion-determined species (species found only in one of the three ecoregions). Species occurrence was coded as 1 (present) or 0 (absent) across 18 simulated sites in three different ecoregions with a fixed proportion of species for each of the above types (7:2:1 ecoregion:random:panmixia) spread across the ecoregions (7:2:1 ecoregion:random:panmixia). These observations were then subject to incidence values changes (1 changed to 0 or 0 changed to 1) at varying proportions across the entire dataset (30% to 44% of data), these proportions produced ecoregion separation on nMDS plots similar to empirical ecological datasets during initial trials. The output of this simulation was visualised using an nMDS ordination of Jaccard dissimilarities (see Fig. SI5.1). A PERMANOVA was used to assess the significance of differences in multivariate location between the three ecoregions. As an ecoregion difference was simulated in the community data a p-value greater than 0.05 indicates a false negative result due to stochasticity.

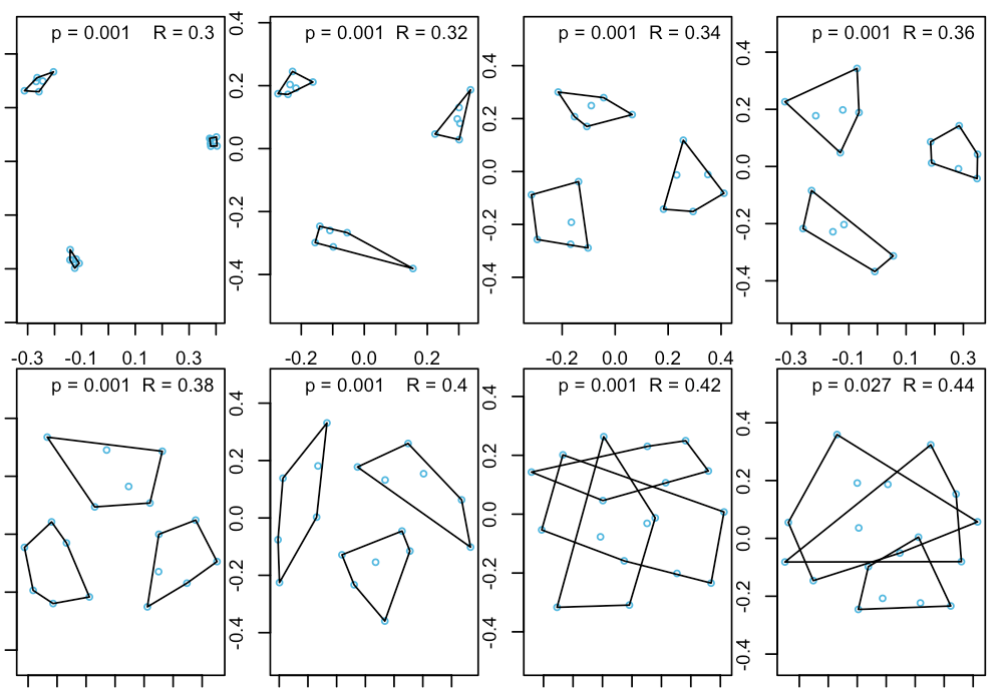

***Figure SI5.1****. Non-metric multidimensional scaling ordinations based on Jaccard dissimilarities of simulated community data. Simulated communities had 18 sites equally spaced among three ecoregions and 1000 species were simulated. A proportion of species incidence records were randomised from 30% (R=0.3) to 44% (R=0.44). Sites are shown in blue and a convex hull is shown for each ecoregion. The p value reported in each simulation corresponds to the PERMANOVA result testing for significant differences in the multivariate spread of ecoregions.*

This simulation was replicated 100 times for 20, 50, 100, 500, 1000 and 2000 species (Fig. SI5.2).

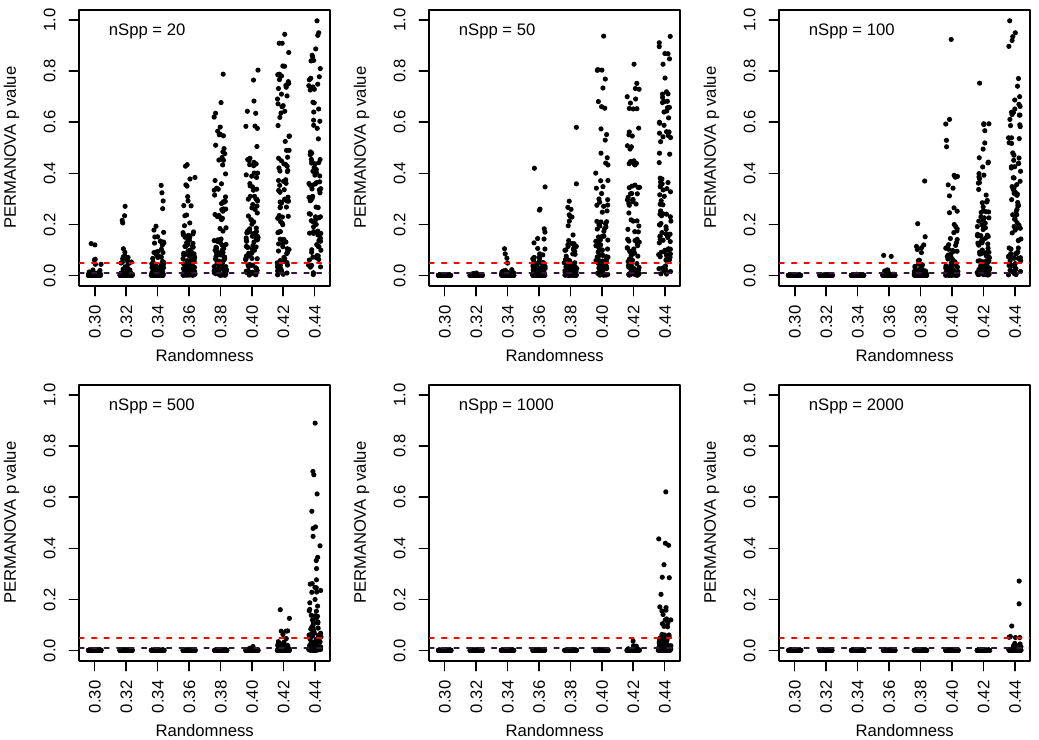

***Figure SI5.2****. Plots showing the significance value (p value) for a PERMANOVA testing multivariate location between three ecoregions in simulated community data. Simulations were conducted for eight levels of randomness values: indicating the proportion of observations changes to an alternative state (present to absent or vice versa). Each plot shows 100 simulations for each level of randomness, with different plots showing effect for an increasing number of simulated species observations (nSpp). The red dashed lines indicate a PERMANOVA p-value of 0.05 and the dark purple dashed line 0.01.*

The ordinations between 0.32 and 0.38 showed the highest similarity to those presented in the main manuscript (Fig. 2). The proportion of false negative simulations was between 0.15-0.71 for 20 species and 0-0 for 2000 species (see Table SI5.1). Between 50 and 100 species the false negative proportion decreased from an average of 0.13 to 0.03, representing a shift from 87% to 97% of simulations finding a significant effect of ecoregions. Therefore, a reasonably high (0.95) chance of seeing a true effect lies between 50 and 100 species.

***Table SI5.1*** *Proportion of false negative results (p >0.05) from a PERMANOVA significance test of ecoregion across 100 simulated datasets. Each dataset was simulated for a given randomness value (0.3-0.44 – shown on left) and number of species (20-2000).*

|  | **Number of Species** | | |  |  |  |
| --- | --- | --- | --- | --- | --- | --- |
| **Randomness** | **20** | **50** | **100** | **500** | **1000** | **2000** |
| **0.3** | 0.04 | 0 | 0 | 0 | 0 | 0 |
| **0.32** | 0.15 | 0 | 0 | 0 | 0 | 0 |
| **0.34** | 0.28 | 0.03 | 0 | 0 | 0 | 0 |
| **0.36** | 0.53 | 0.21 | 0.02 | 0 | 0 | 0 |
| **0.38** | 0.71 | 0.28 | 0.1 | 0 | 0 | 0 |
| **0.4** | 0.8 | 0.72 | 0.39 | 0 | 0 | 0 |
| **0.42** | 0.87 | 0.69 | 0.71 | 0.06 | 0 | 0 |
| **0.44** | 0.92 | 0.9 | 0.86 | 0.56 | 0.26 | 0.07 |

### Supplementary Information 5

*Distance-decay relationship analyses*

**Analysis by taxonomic group**

Distance-decay slopes for all observations showed an exponential decrease in compositional similarity as the distance between sites increased (Fig. SI4.1, see below). Regression models of log10 transformed compositional similarity indicated that this slope was statistically significant in all cases (p <0.001 for all markers & taxonomic groups, full model output below). In the metazoan datasets (both COI & 18S) the model showed a significant difference in the slope between artificial and natural sites (COI - p = 0.0002; 18S - p = 0.0019, full model outputs below). No statistically significant differences were found between site types in the protist or bacteria datasets.

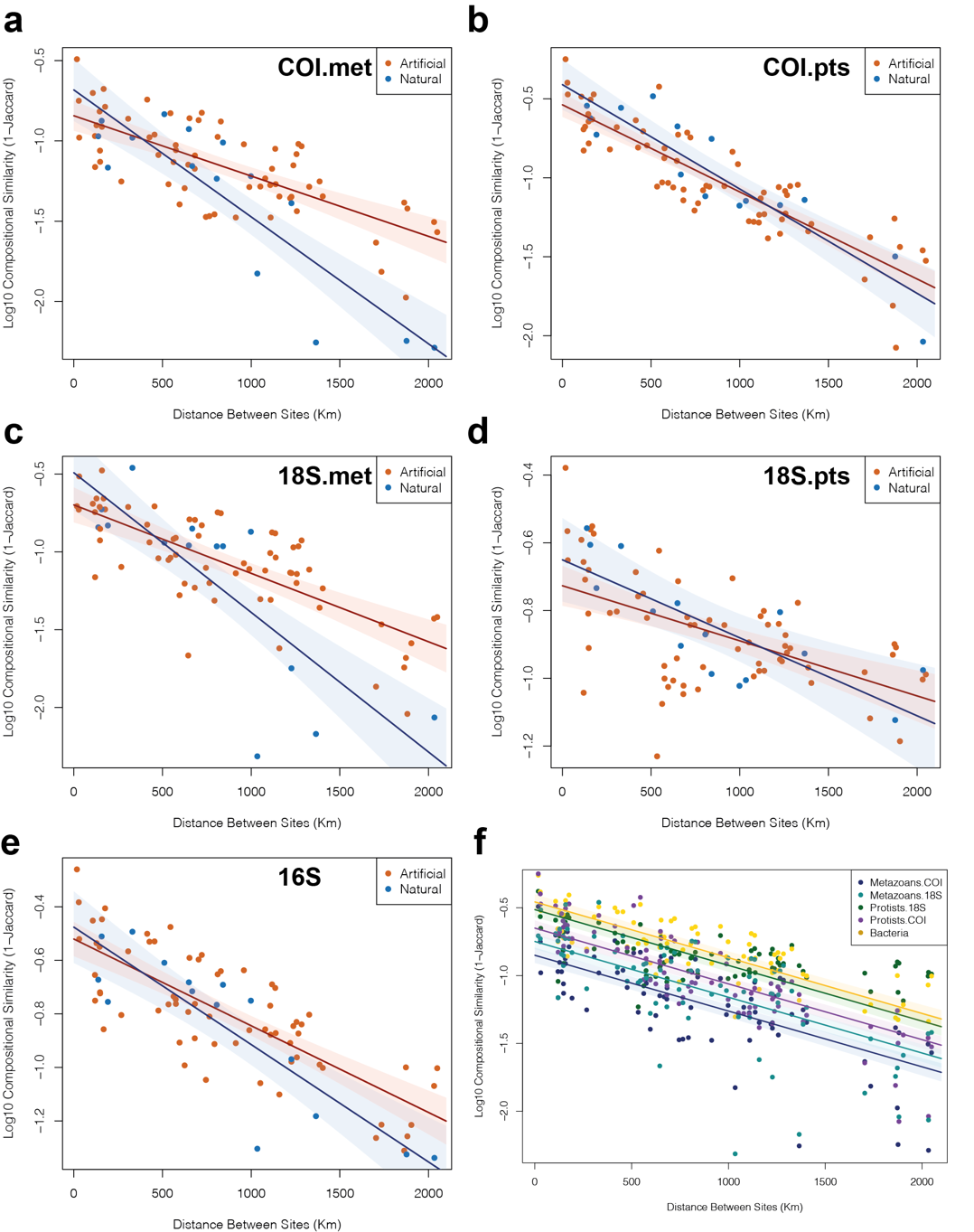

**Figure SI4.1**. *Plots showing distance between sites and community dissimilarity measured by environmental DNA metabarcoding across South Africa. Logarithmically (base 10) transformed compositional similarity against distance is shown for* ***a*** *COI metazoans,* ***b*** *COI protists,* ***c*** *18S metazoans* ***d*** *18S protists and* ***e*** *16S data with comparisons between artificial sites coloured red and natural sites coloured blue. 95% confidence intervals from the least-squares linear regression models are shown as light shaded areas around each regression slope.* ***f*** *All distance decay data overlayed.*

**Table SI4.1** *Least-squares regression model for Distance-Decay relationships between different site types measured by eDNA metabarcoding for coastal sites across South Africa. Models are presented for three genes; COI – cytochrome c oxidase subunit I; 18S -nuclear small subunit ribosomal DNA, 16S – prokaryotic small subunit ribosomal DNA.*

| COI metazoan | |  |  |  |
| --- | --- | --- | --- | --- |
| F_(3,76)_=47.73 |  |  |  |  |
| R2 =0.6396 | Estimate | Std.Error | t value | P value |
| (Intercept) | -8.43E-01 | 4.71E-02 | -17.922 | **<2e-16** |
| distance | -3.76E-04 | 4.77E-05 | -7.88 | **1.87E-11** |
| site.typeNatural | 1.61E-01 | 1.08E-01 | 1.488 | 0.140856 |
| distance:site.typeNatural | -4.15E-04 | 1.06E-04 | -3.905 | **0.000202** |
| COI protista |  |  |  |  |
| F_(3,77)_=94.5 |  |  |  |  |
| R2 =0.7781 |  |  |  |  |
| (Intercept) | -5.38E-01 | 3.82E-02 | -14.068 | **<2e-16** |
| distance | -5.52E-04 | 3.79E-05 | -14.534 | **<2e-16** |
| site.typeNatural | 1.28E-01 | 8.86E-02 | 1.441 | 0.154 |
| distance:site.typeNatural | -1.09E-04 | 8.67E-05 | -1.261 | 0.211 |
| 18S metazoan | |  |  |  |
| F_(3,76)_=37.22 |  |  |  |  |
| R2 =0.579 | Estimate | Std.Error | t value | P value |
| (Intercept) | -6.98E-01 | 5.63E-02 | -12.394 | **<2e-16** |
| distance | -4.40E-04 | 5.59E-05 | -7.867 | **1.98E-11** |
| site.typeNatural | 2.07E-01 | 1.35E-01 | 1.534 | 0.12916 |
| distance:site.typeNatural | -4.58E-04 | 1.43E-04 | -3.213 | **0.00193** |
| 18S protista |  |  |  |  |
| F_(3,77)_=15.09 |  |  |  |  |
| R2 =0.3458 |  |  |  |  |
| (Intercept) | -7.27E-01 | 2.97E-02 | -24.504 | **<2e-16** |
| distance | -1.63E-04 | 2.94E-05 | -5.528 | **4.25E-07** |
| site.typeNatural | 7.65E-02 | 6.87E-02 | 1.114 | 0.269 |
| distance:site.typeNatural | -6.76E-05 | 6.72E-05 | -1.006 | 0.318 |
| 16S |  |  |  |  |
| F_(3,77)_=25.82 |  |  |  |  |
| R2 =0.4821 |  |  |  |  |
| (Intercept) | -1.52E-01 | 9.04E-03 | -16.826 | **<2e-16** |
| distance | 7.16E-05 | 8.97E-06 | 7.978 | **1.12E-11** |
| site.typeNatural | 1.22E-02 | 2.09E-02 | 0.583 | 0.562 |
| distance:site.typeNatural | -4.68E-06 | 2.05E-05 | -0.229 | 0.82 |

**Analysis by gene region**

We analysed the distance-decay slopes for all observations and these showed an exponential decrease in compositional similarity as the distance between sites increased (Fig. SI4.2 below). Regression models of log10 transformed compositional similarity indicated that this slope was statistically significant in all cases (P<0.005 for all markers, full model output below). In the COI data the model showed a significant difference in both the slope and intercept between artificial and natural sites (F_3,77_=77.92, additive p=0.018, interactive p=0.004, full model below). No statistically significant difference was found between site types in the 18S or 16S data. As shown in Fig. 3b-d, there was much greater variance in the residuals for the 18S and 16S data compared to the COI data and the R^2^ values from the models support these observations (COI - R^2^=0.730, 18S - R^2^=0.224, 16S - R^2^=0.473).

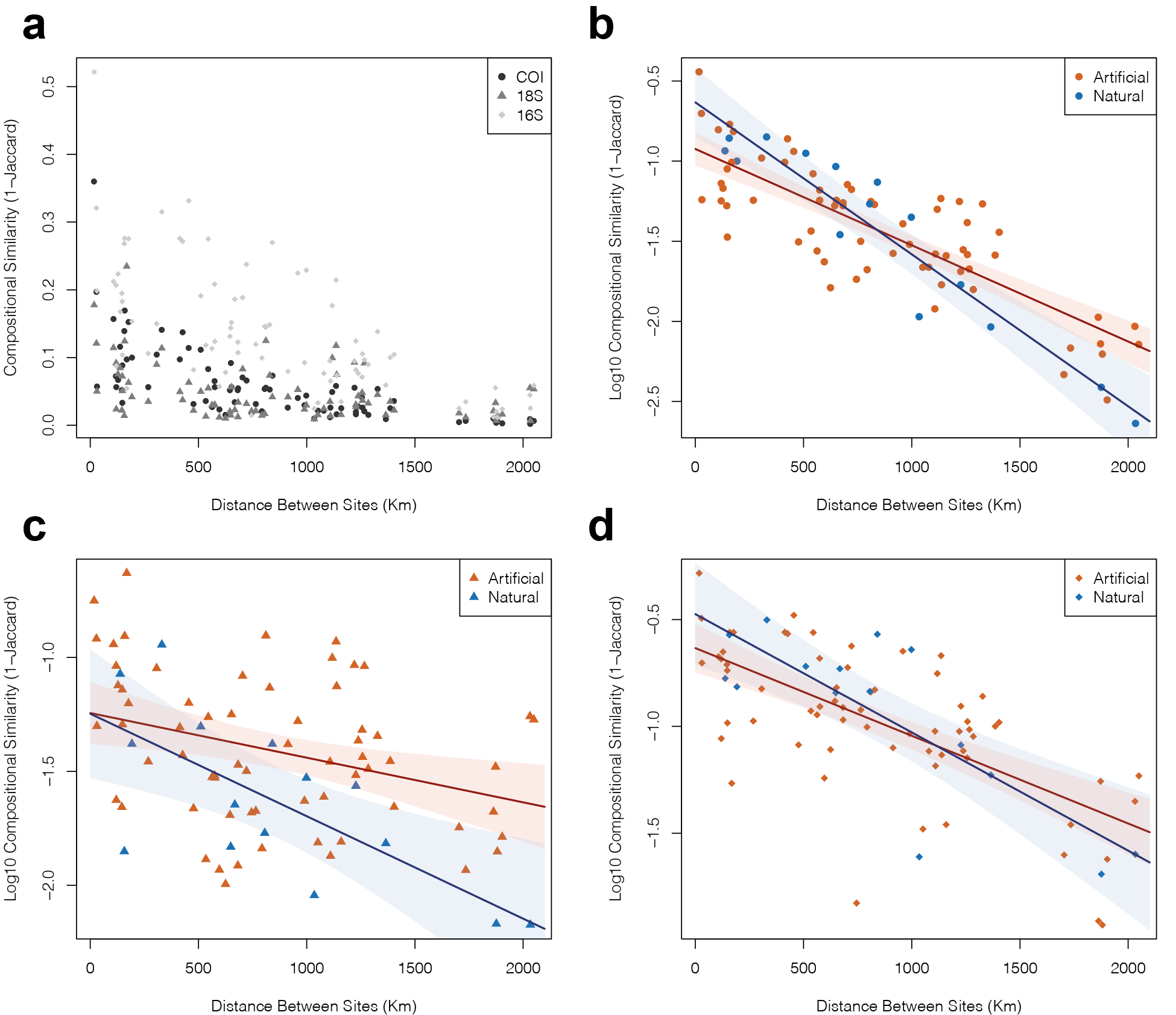

***Figure SI4.2****. Plots showing distance between sites and community dissimilarity measured by environmental DNA metabarcoding across South Africa.* ***a*** *Untransformed data for all gene regions. Logarithmically (base 10) transformed compositional similarity against distance is shown for* ***b*** *COI,* ***c*** *18S and* ***d*** *16S data with comparisons between artificial sites coloured red and natural sites coloured blue. 95% confidence intervals from the least-squares linear regression models are shown as light shaded areas around each regression slope.*

**Table SI4.2** *Least-squares regression model for Distance-Decay relationships between different site types measured by eDNA metabarcoding for coastal sites across South Africa. Models are presented for three genes; COI – cytochrome c oxidase subunit I; 18S -nuclear small subunit ribosomal DNA, 16S – prokaryotic small subunit ribosomal DNA.*

| COI |  |  |  |  |
| --- | --- | --- | --- | --- |
| F_(3,77)_=77.92 |  |  |  |  |
| R2 =0.7295 | Estimate | Std.Error | t value | P value |
| (Intercept) | -9.23E-01 | 5.17E-02 | -17.849 | **<2.00E-16** |
| distance | -6.02E-04 | 5.13E-05 | -11.726 | **<2.00E-16** |
| site.typeNatural | 2.89E-01 | 1.20E-01 | 2.415 | **0.01812** |
| distance:site.typeNatural | -3.48E-04 | 1.17E-04 | -2.967 | **0.00401** |
| 18S |  |  |  |  |
| F_(3,77)_=8.679 |  |  |  |  |
| R2 =0.2236 |  |  |  |  |
| (Intercept) | -1.2434844 | 0.0677062 | -18.366 | **<2e-16** |
| distance | -0.0001962 | 0.0000672 | -2.919 | **0.0046** |
| site.typeNatural | -0.0015438 | 0.1568279 | -0.01 | 0.9922 |
| distance:site.typeNatural | -0.0002571 | 0.0001535 | -1.675 | 0.0979 |
| 16S |  |  |  |  |
| F_(3,77)_=24.94 |  |  |  |  |
| R2 =0.4731 |  |  |  |  |
| (Intercept) | -6.33E-01 | 5.72E-02 | -11.076 | **<2.00E-16** |
| distance | -4.10E-04 | 5.68E-05 | -7.219 | **3.19E-10** |
| site.typeNatural | 1.58E-01 | 1.33E-01 | 1.195 | 0.236 |
| distance:site.typeNatural | -1.44E-04 | 1.30E-04 | -1.107 | 0.272 |
